# Systematic Evaluation of Machine Learning Algorithms for Neuroanatomically-Based Age Prediction in Youth

**DOI:** 10.1101/2021.11.24.469888

**Authors:** Amirhossein Modabbernia, Heather C. Whalley, David C. Glahn, Paul M. Thompson, Rene S. Kahn, Sophia Frangou

**Author notes:** Corresponding author: Sophia Frangou, Department of Psychiatry, Icahn School of Medicine at Mount Sinai, 1 Gustave L Levy Place, New York NY 10029.

## Abstract

Application of machine learning algorithms to structural magnetic resonance imaging (sMRI) data has yielded behaviorally meaningful estimates of the biological age of the brain (brain-age). The choice of the machine learning approach in estimating brain-age in children and adolescents is important because age-related brain changes in these age-groups are dynamic. However, the comparative performance of the multiple machine learning algorithms available has not been systematically appraised. To address this gap, the present study evaluated the accuracy (Mean Absolute Error; MAE) and computational efficiency of 21 machine learning algorithms using sMRI data from 2,105 typically developing individuals aged 5 to 22 years from five cohorts. The trained models were then tested in two independent holdout datasets, comprising 4,078 pre-adolescents aged 9-10 years and another sample of 594 individuals aged 5-21 years. The algorithms encompassed parametric and nonparametric, Bayesian, linear and nonlinear, tree-based, and kernel-based models. Sensitivity analyses were performed for parcellation scheme, number of neuroimaging input features, number of cross-validation folds, number of extreme outliers, and sample size. The best performing algorithms were Extreme Gradient Boosting (MAE of 1.49), Random Forest Regression (MAE of 1.58) and Support Vector Regression with Radial Basis Function Kernel (MAE of 1.64) which had acceptable and comparable computational efficiency. Findings of the present study could be used as a guide for optimizing methodology when quantifying age-related changes in youth.

## 1. Introduction

Brain development involves highly organized multistep processes (Tau and Peterson, 2010) that lead to the emergence of adult levels of cognitive and behavioral competency (Paus, 2005; Spear, 2000). Brain development involves numerous cellular and noncellular events (Tau and Peterson, 2010), which are below the resolution of magnetic resonance imaging (MRI) but underpin morphological changes in brain organization that can be captured using structural MRI (sMRI) techniques. Multiple studies have shown that the volume of subcortical structures typically peaks in late childhood and adolescence and decreases thereafter (Dima et al., 2022; Raznahan et al., 2014). Cortical thickness shows a steep reduction in late childhood and adolescence that continues at a slower rate throughout adult life (Frangou et al., 2022; Wierenga et al., 2022). Cortical surface area expands during childhood and most of adolescence showing gradual decrements thereafter (Fjell et al., 2015; Tamnes et al., 2017).

These age-related changes demonstrate marked inter-regional and inter-individual variation (Mills et al., 2021; Wierenga et al., 2022).

Machine learning (ML) algorithms applied to sMRI data can harness the multidimensional nature of age-related brain changes at the individual-level to predict age, as a proxy for the biological age of the brain (i.e., brain-age). The difference between brain-age and chronological age is referred to here as brain-age-gap-estimation (BrainAGE)(Franke and Gaser, 2019), which is equivalent to terms such as brain-predicted-age-difference (brainPAD) (Luna et al., 2021), brain-age-gap (BAG) (Anaturk et al., 2021) and brain-age delta(Beheshti et al., 2019) used in other studies. In adults, higher brain-age relative to chronological age (i.e., higher BrainAGE) has been associated with adverse physical (Cole et al., 2018), cognitive (Anaturk et al., 2021; Boyle et al., 2021; Elliott et al., 2019) and mental health phenotypes (Kaufmann et al., 2019; Lee et al., 2021). By contrast, in children and adolescents higher BrainAGE has been associated with better cognitive test performance (Boyle et al., 2021; Erus et al., 2015; Luna et al., 2021) while associations with clinical phenotypes show a more complex pattern which may depend on the nature of the phenotype and/or the developmental stage of the sample (Chung et al., 2018; Luna et al., 2021). These findings underscore the importance of accuracy in brain-based age-prediction in youth, as childhood and adolescence are arguably periods of the most dynamic brain re-organization.

Therefore, the current study focuses exclusively on the evaluation of the methods used to compute brain-age from sMRI data as a foundation for guiding study design into its functional significance. We have previously shown that age prediction from sMRI data in adults is influenced by the choice of algorithm (Lee et al., 2021). Here addressed this knowledge gap in youth because with few exceptions (Ball et al., 2021; Brouwer et al., 2021; Lee et al., 2021; Luna et al., 2021), studies on brain-age prediction in this population have typically employed a single ML algorithm, most commonly relevance vector regression (RVR), Gaussian process regression (GPR), and support vector regression (SVR) (Cole et al., 2018; Franke et al., 2012; Franke et al., 2010; Gaser et al., 2013; Liem et al., 2017; Valizadeh et al., 2017). Here we systematically evaluated the performance of 21 ML algorithms applied to sMRI data from youth from five different cohorts and then tested their performance in two independent samples. The algorithms encompassed parametric and nonparametric, Bayesian, linear and nonlinear, tree-based, and kernel-based models. These algorithms were selected to include those that are commonly used in brain-age prediction studies as well representative examples of a range of algorithms that provide reasonable and potentially better alternatives. We evaluated the ML methods for accuracy and for their sensitivity to key parameters known to affect model performance pertaining to parcellation scheme, number of neuroimaging input features (Valizadeh et al., 2017), number of cross-validation folds, sample size (by resampling the available data), and number of extreme outliers. Ourprediction was that nonlinear kernel-based and ensemble algorithms would outperform other algorithms because they are theoretically better at handling colinear data and non-linear relationships with age and, in the case of ensemble algorithms, they improve predictive performance by aggregating results from multiple nodes. Collectively, these analyses may assist in optimizing the design of future investigations on brain predicted age in youth.

## 2. Method

### 2.1 Samples

We used T1-weighted scans from six separate cohorts: Autism Brain Imaging Data Exchange (ABIDE) (Di Martino et al., 2017; Di Martino et al., 2014); ABIDE II (Di Martino et al., 2017); ADHD-200 (Consortium, 2012); Human Connectome Project Development (HCP-D) (Harms et al., 2018); Child Mind Institute (CMI) (Alexander et al., 2017), Adolescent Brain Cognitive Development (ABCD) (Garavan et al., 2018), Pediatric Imaging, Neurocognition, and Genetics (PING) Data Repository (Jernigan et al., 2016) (details of the cohorts in the Supplemental Material). Data collection for these cohorts was conducted at multiple independent sites located in eight countries: (USA, Germany, Ireland, Belgium, the Netherlands, Switzerland, China, and France). Only psychiatrically healthy participants with high-quality anatomical brain scans from each cohort were included (Supplemental Material and Supplemental Table 1). Data from five cohorts (total n=2,105; 41% female, age-range: age-range:9-10years) (Supplemental Figure 1) were used to train the ML algorithms (training set) while data from the ABCD sample (n=4,078; 52% female; age range: 9-10 years) and the PING sample (n=594; female=49.6%; age-range 5-21 years) comprised the independent hold-out test-sets.

### 2.2 Image processing

Across all cohorts, more than 98% of the participants were scanned using 3-T MRI machines; Siemens Prisma and Trio Tim scanners were each used for 31% of the participants of the total training sample (Supplemental Table 1). The T1-weighted images were downloaded from the respective cohort repositories and processed at the Icahn School of Medicine at Mount Sinai (ISMMS) using identical pipelines. Image processing was implemented using standard pipelines in the FreeSurfer 7.1.0 software to generate cortical parcels based on the Schaefer scheme (Schaefer et al., 2018) by projecting the parcellation onto individual surface space (https://github.com/ThomasYeoLab/CBIG/tree/master/stable_projects/brain_parcellation/Schaefer2018_LocalGlobal/Parcellations/project_to_individual) and using the *mri_anatomical_stats* function to extract cortical values. We used the 400-parcel resolution (i.e., 400 cortical thickness and 400 cortical surface area values) (Supplemental Figure 2) in the main analyses. The 400-parcellation scheme has been shown to have good stability, signal to noise ratio, and performance in different contexts, and correspondence to histology (Bryce et al., 2021; Valk et al., 2020). Participants with missing values on any parcellations were excluded, because it was assumed that the image quality was compromised; participants in the training dataset only were also excluded if more than 5% of their parcellation features had extreme values (details in Supplemental Material). We did not exclude participants based on outlier values in the hold-out test sets but instead studied the effect of outliers on model performance.

### 2.3 Algorithms for brain-based age prediction

We used the *caret* package (version 6.0.84) in R (version 3.5.3) to conduct the ML analyses because it interfaces with multiple ML packages and standardizes data preprocessing, model training and testing. Several of the regression algorithms evaluated can be extended to accommodate non-linear associations using kernel functions. A kernel function transforms the original non-linear data into a higher-dimensional space in which they can become linearly separable. The kernelized models evaluated here incorporated polynomial and radial basis function (RBF) kernels. The former adds features using the polynomial combinations of the original data up to a specified degree and the latter adds features using the distance of the original data from specified reference values. Below we describe the 15 base models, six of which have non-linear kernelized variations; together, they amount to 21 different algorithms:

1. **Generalized linear model:** This is a standard algorithm for regression that minimizes the sum of squared errors between the observed variables and predicted outcomes. Models have no tuning parameters and were implemented using the **“**glm” function.
2. **Bayesian general linear model** (Gelman et al., 2008): This a linear regression model in which the outcome variable and the model parameters are assumed to be drawn from a probability distribution; it therefore provides estimates of model uncertainty. Models have no tuning parameters and were implemented using the “bayesglm” function.
3. **Gaussian Processes Regression** (Williams and Barber, 1998): This is a regression model that follows Bayesian principles. The covariance function here was defined by using either a linear function, or a polynomial or a RBF kernel as a prior. The polynomial kernels were tuned using degree and scale and the RBF kernels were tuned using the *sigma* parameter (the inverse kernel width parameter). Models were implemented using “gaussprRadial”, “gaussprLinear”, and “gaussprPoly”.
4. **Independent Component Regression** (Shao et al., 2006): This is a linear regression model in which components from a prior independent component analysis are used as the explanatory variables. The number of components was tuned, and the models were implemented using the “icr” function.
5. **Principal Component Regression**: This is a linear regression model in which components from a prior principal component analysis are used as the explanatory variables. The number of components was tuned, and the models were implemented using the “pcr” function.
6. **Kernel Partial Least Squares Regression** (Dayal and MacGregor, 1997): This is an extension of the partial least squares (PLS) regression which creates components by using the correlations between explanatory variables and outcome variables. The kernelized version used here (K-PLS) maps the data vector from the sample space to a higher-dimensional, Euclidean space; models were tuned for the number of components and implemented using the “kernelpls” function.
7. **Sparse Partial Least Squares Regression (SPLS)**(Chun and Keleş, 2010): This is a different extension of PLS that reduces the number of explanatory variables (sparsity) through a least absolute shrinkage and selection operator (LASSO) approach. The models were tuned for the number of components, and *eta* (the sparsity parameter), and were implemented using the “spls” function.
8. **Quantile Regression with least absolute shrinkage and selection operator (LASSO) Penalty** (Wu and Liu, 2009): This algorithm models the relationship between explanatory variables and specific percentiles (or “quantiles”) of the outcome variable; in this variation, sparsity was introduced through the LASSO approach. The number of selected variables was tuned, and models were implemented with the “rqlasso” function.
9. **Elastic Net Regression** (Zou and Hastie, 2005): This is a linear regression that adds two penalties, LASSO regression (L1-norm) and ridge regression (L2-norm), in the loss function to encourage simpler models and avoid overfitting. Models were tuned for *lambda* (weight decay) and fraction of the full solution (equivalent to ordinary least squares) and were implemented using the “enet” function.
10. **Boosted Generalized Additive Model** (Bühlmann and Yu, 2003): This generalized additive model is fitted using a gradient-based boosting algorithm based on penalized B-splines. Overfitting was reduced by pruning the number of iterations using the optimal value of the Akaike Information Criterion. Models were implemented using the “gamboost” function.
11. **Random Forest Regression** (Breiman, 2001): This an ensemble machine learning method, which involves construction of multiple decision trees (i.e., forests) via bootstrap (bagging) and aggregates the predictions from these multiple trees to reduce the variance and improve the robustness and precision of the results. Models were implemented using the “rf” function and were tuned with regard to the number of trees.
12. **Support Vector Regression** (Cortes and Vapnik, 1995): Support Vector Regression (SVR) is characterized by the use of kernels, sparsity, and control of the margin of tolerance (epsilon; ε) and the number of support vectors (Awad and Khanna, 2015). It identifies a symmetrical ε-insensitive region, called the ε-tube, which approaches the loss function as an optimization problem; the ε-value determines the width of the tube and maximization of the “flatness” aims to ensure that it contains most of the values in the training sample. Here flatness maximization was subject to the L2-norm penalty. In addition to the linear kernel, we also tested a version with polynomial and RBF kernels. The corresponding functions were “svmLinear3”, “svmPoly” and “svmRadial”. The regularization parameter (C) was used to optimize all models, while scale and degree were also considered in polynomial models and *sigma* for RBF models.
13. **Relevance Vector Regression** (Tipping, 2001): Relevance Vector Regression (RVR) is an extension of SVR embedded in a Bayesian framework. Its characteristic feature is that it imposes an explicit zero-mean Gaussian prior on the model parameters leading to a vector of independent hyperparameters that reduces the dataset. The behavior of the RVR is controlled by the type of kernel, which has to be chosen, while all other parameters are automatically estimated by the learning procedure itself. Here we used a linear, polynomial, or RBF kernel implemented with functions “rvmLinear” “rvmPoly” and “rvmRadial” respectively. The latter two kernels require tuning for scale and degree (polynomial) and for sigma (RBF).
14. **Bayesian Regularized Neural Networks** (Perez-Rodriguez et al., 2013): This is a version of the feedforward artificial neural network (ANN) architecture, in which robustness is improved through Bayesian regularization of the ANN parameters. The model includes two layers: the input layer - consisting of independent variables - and the hidden layer of *S* number of neurons. Models were implemented using the “brnn” function and tuned for the number of neurons.
15. **Extreme Gradient Boosting** (Chen and Guestrin, 2016): Extreme Gradient Boosting (XGBoost) is an ensemble decision-tree based gradient boosting algorithm that allows for modeling complex nonlinear relationships and interactions. The algorithm optimizes model performance through parallel (simultaneous) processing, regularization, tree pruning, optimal split (through a weighted quantile sketch algorithm), automatic missing data handling and built-in cross-validation. Tuning parameters involved the number of boosting iterations; maximum tree depth; eta (shrinkage parameter); gamma (minimum loss reduction); subsample ratio of columns; minimum sum of instance weights, and column subsample percentage. Models were implemented using “xgbTree” function.

For clarity we refer to each algorithm by the name of the specific function used for its implementation.

Computational efficiency for each algorithm was assessed by recording the total Central Processing Unit (CPU) time, and the average and maximum memory usage. All models were run on the ISMMS high-performance computing cluster.

Several analytic steps were common to all algorithms. As there are known sex differences in the rate of age-related changes (Brouwer et al., 2021; Wierenga et al., 2019; Wierenga et al., 2022), models were separately trained for males and females. Hyperparameter tuning (when required) was performed in the combined training set (n=2,105), using a grid search in a five-fold cross-validation scheme across five repeats. In each cross-validation 80% of the training sample was used to train the model and 20% was used to test the model parameters. Subsequently, the model was re-trained on the whole training dataset using the optimal hyperparameters identified through cross-validation. Finally, the generalizability of the model was tested in two hold-out datasets (ABCD n=4,078 and PING n=594).

The primary accuracy measure for each algorithm was the Mean Absolute Error (MAE) which represents the absolute difference between the neuroimaging-predicted age and the chronological age. For each algorithm, the abbreviation MAE_T_ refers to values obtained in the hold-out test dataset and MAE_cv_ refers to the mean cross-validation value in the training dataset. We also report two other commonly used accuracy measures: the Root Mean Square Error (RMSE), which is the standard deviation of the prediction errors, and the correlation between predicted and actual age. This correlation coefficient was not calculated in the ABCD data because of the narrow age range (less than 2 years). Based on these criteria we identified the three best performing algorithms which we evaluate further in the subsequent sections.

### 2.4 Calculating BrainAGE and corrected BrainAGE for the three best performing algorithms

BrainAGE in each individual was calculated by subtracting the chronological age from the age predicted by each of the three best performing algorithms. Positive BrainAGE values indicate an older than expected brain-age for the given chronological age, and the opposite is the case for negative BrainAGE values. BrainAGE is typically overestimated in younger individuals and underestimated in older individuals. To counter this bias, multiple methods have been proposed (Beheshti et al., 2019; Cole et al., 2018). Here we used a robust approach introduced by Beheshti and colleagues (Beheshti et al., 2019), which relies on the slope (α) and intercept (β) of a linear regression model of BrainAGE against chronological age in the training set. This way an offset is calculated (as αΩ+β) and then subtracted from the estimated brain-age to yield a bias-free BrainAGE (Beheshti et al., 2019), hereafter referred to as “corrected BrainAGE” (BrainAGE_corr_).

### 2.5 Quantifying feature importance for age prediction in the three best performing algorithms

Estimates of the contribution of individual neuroimaging features to age prediction in each of the three best performing algorithms were obtained using Shapley Values (SV) implemented via the *fastshap* package Version 0.0.5 (Greenwell and Greenwell, 2020) in R. SVs derive from the cooperative game theory,(Lundberg and Lee, 2017), they accommodate non-linearity and have properties that make their interpretation intuitive. For example, the sum of all the SVs of a model is equal to the accuracy of the model and features with the same SV contribute equally to the model.

### 2.6 Sensitivity and supplemental analyses in the three best performing algorithms

To test the effect of sex on model generalizability, we applied the parameters trained on one sex to the other and compared differences in BrainAGE using a two-sample Student’s t test. Sensitivity analyses focused on the parcellation scheme, number of input features, sample size and number of repeats and cross-validation folds and outliers. Accordingly, we repeated the analyses using features from (a) the Desikan-Killiany (DK) atlas (n of features=136); (b) the DK and subcortical Aseg atlas in FreeSurfer combined (n of features=157); (c) the 400-parcel Schaefer atlas with Aseg atlas (n of features=821); and (d) the 1000-parcel Schaefer atlas (n of features 2000). To test effect of sample size, the training dataset was randomly resampled with replacement in increments of 100, from 100-1500 (20 times each). Additionally, we conducted the same analyses using 10 repeats and 10 cross-validation folds. Finally, we tested the effect of number of extreme outliers (potential indicators of low-quality segmentation) on the model performance, by calculating the Spearman’s correlation coefficient between the number of outliers and absolute error value among the subjects in the hold-out test sets. An outlier was defined as 3 median absolute deviation above or below the median for each brain region.

## 3. Results

### 3.1 Algorithm performance for age prediction

Linear algorithms, with the exception of Elastic Net Regression, performed poorly while the XGBoost, RF regression and SVR with RBF kernel emerged as the three top performing models in males and females in cross-validation (Supplementary Table 2) and in the hold-out datasets (i.e., ABCD and PING) (Figure 1, Table 1). Nevertheless, the median correlation coefficient between age predicted by the 21 different algorithms was 0.92 for males and 0.94 for females (Figure 2). The wider age-range of the PING dataset enabled examination of the association between observed and predicted age as predicted from the different algorithms which had a median correlation coefficient of 0.84 for males and 0.86 for females (Table 1, Figure 3, Supplemental Figure 3).

**Table 1.**
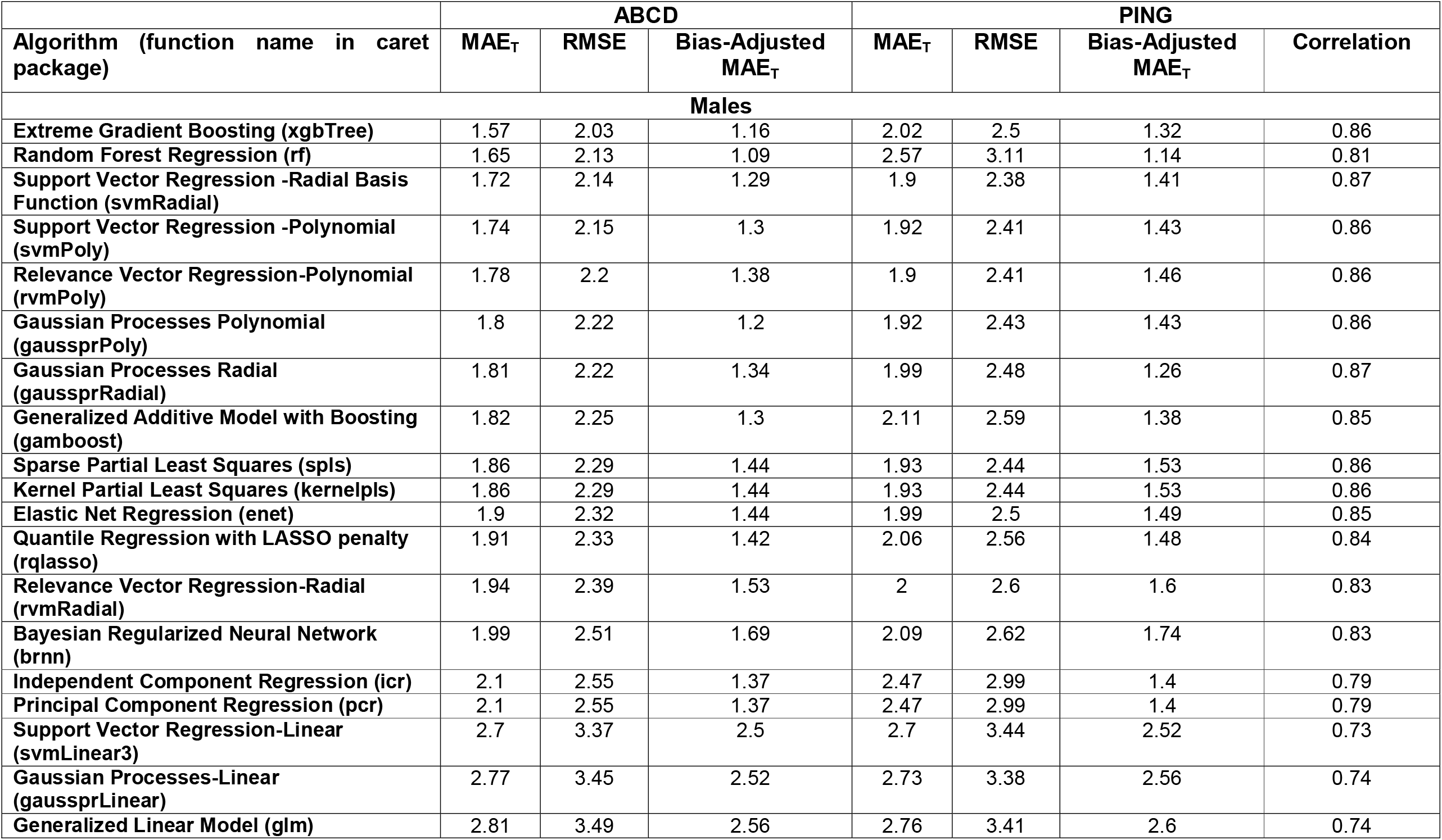
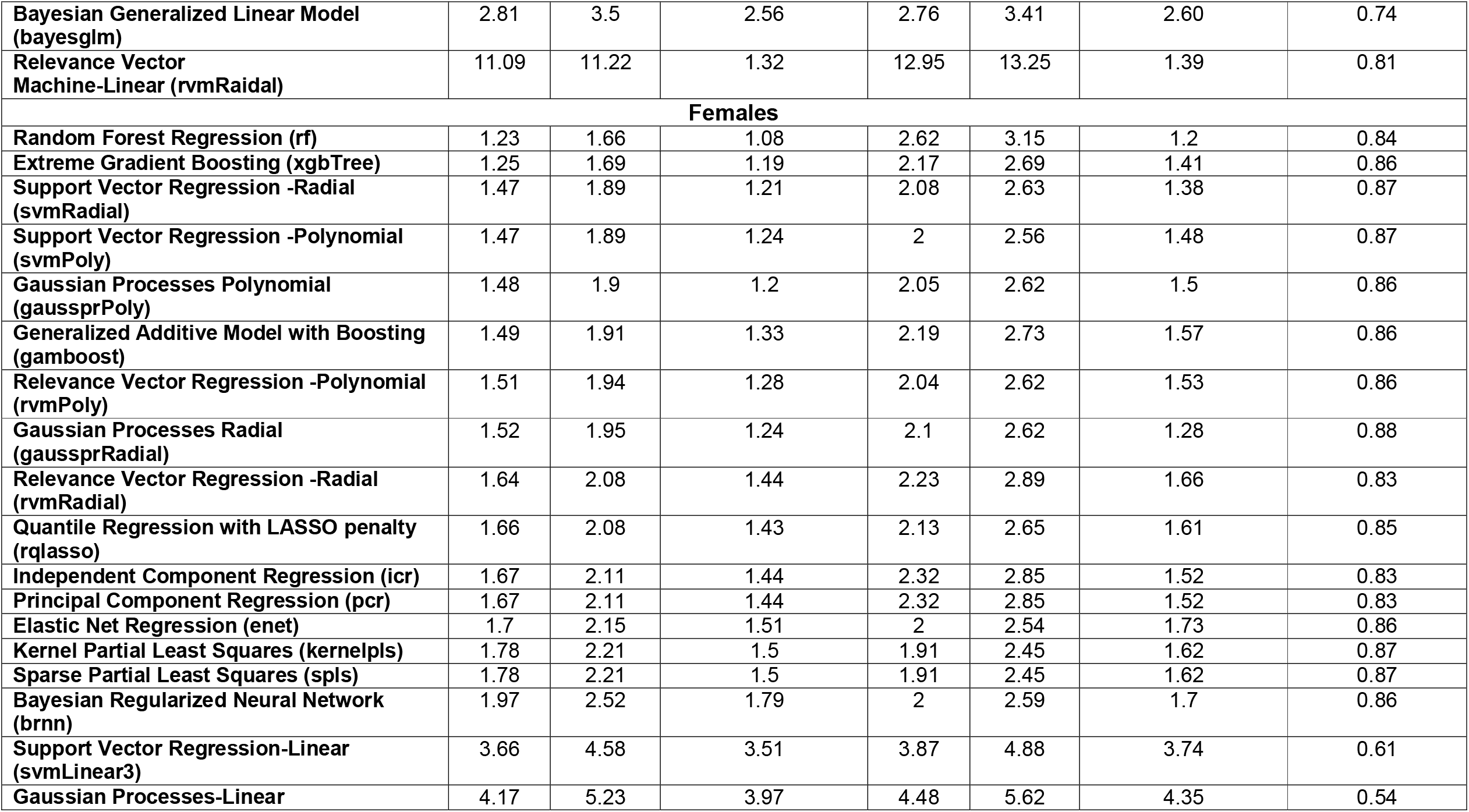
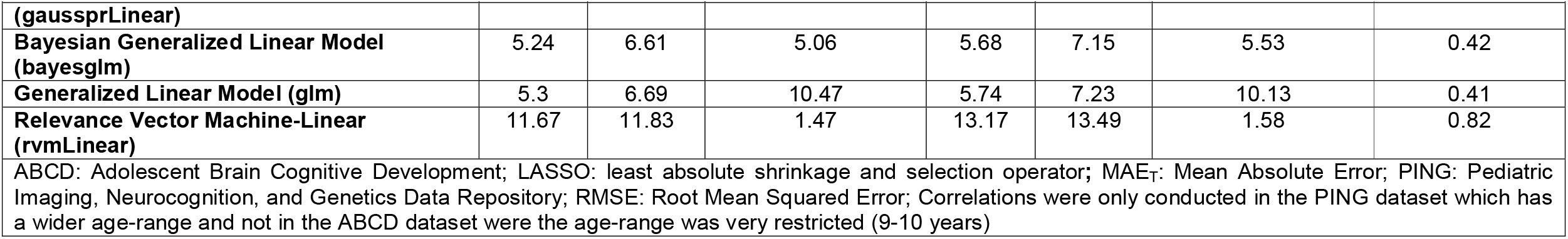
Algorithm Performance in the Hold-out Sets.

**Figure 1.**
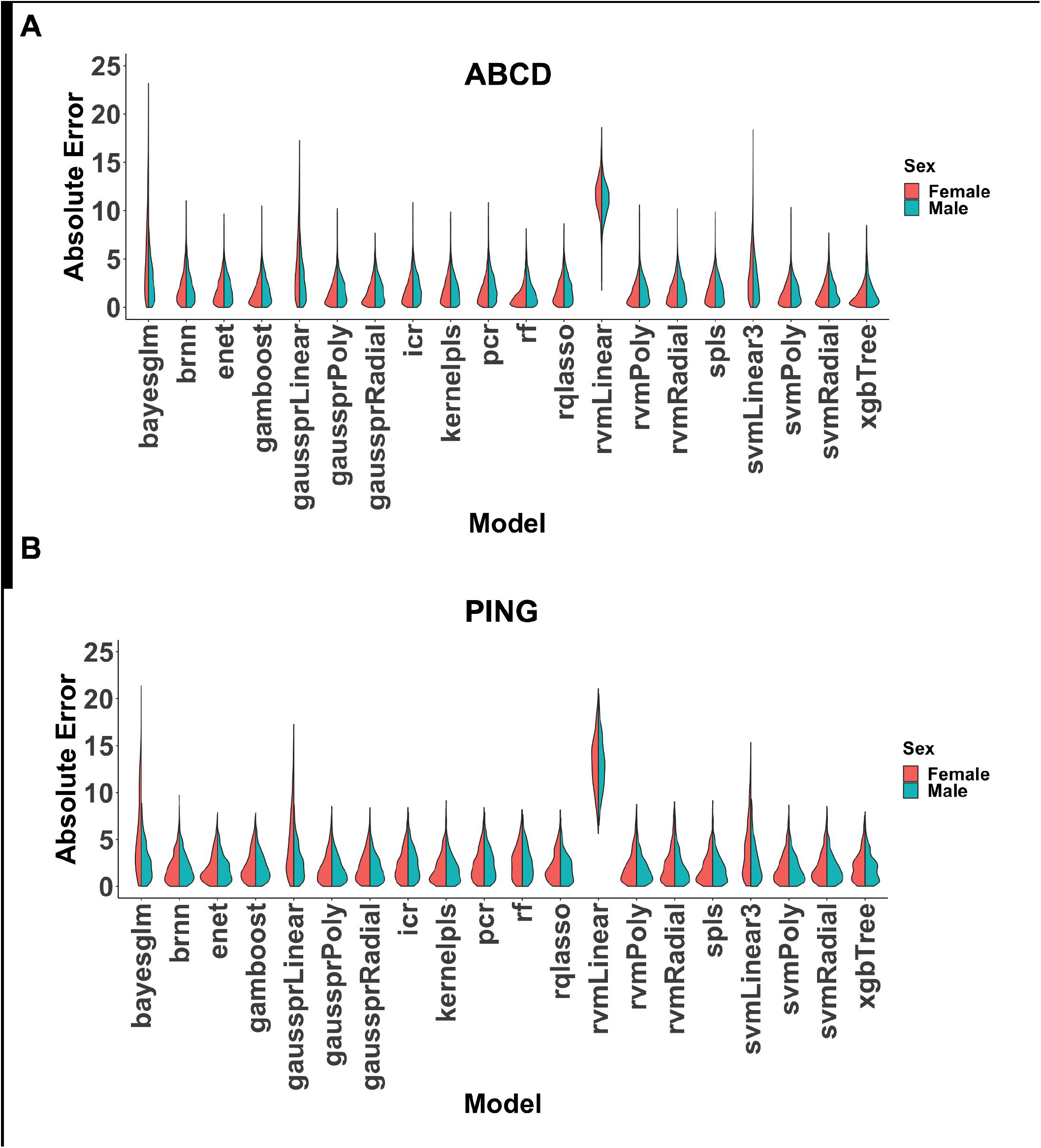
Absolute Error of the 21 Algorithms Evaluated. The figure presents the model performance in males and females in the hold-out test sets: the Adolescent Brain Cognitive Development (ABCD) study (Panel A) and the Pediatric Imaging, Neurocognition, and Genetics Data Repository (PING) (Panel B). The different algorithms are referenced by the function used for their implementation. Bayesian Generalized Linear Model (bayesglm); Bayesian Regularized Neural Network (brnn); Elastic Net Regression (enet);Generalized Additive Model with Boosting (gamboost);: Gaussian Processes Regression Linear (gaussprLinear); Gaussian Processes Regression Polynomial (gaussprPoly); Gaussian Processes Regression Radial (gaussprRadial); Generalized Linear Model (glm); Independent Component Regression (icr); Kernel Partial Least Squares (kernelpls); Principal Component Regression (pcr); Random Forest (rf); Quantile Regression with LASSO penalty (rqlasso); Relevance Vector Machine-Linear (rvmLinear3); Relevance Vector Machine-Polynomial (rvmPoly); rvmRadial: Relevance Vector Machine-Radial; Sparse Partial Least Squares (spls); Support Vector Regression-Linear (svmeLinear3); Support Vector Regression-Polynomial (svmPoly); Support Vector Regression-Radial (svmRadial); Extreme Gradient Boosting (xgbTree)

**Figure 2.**
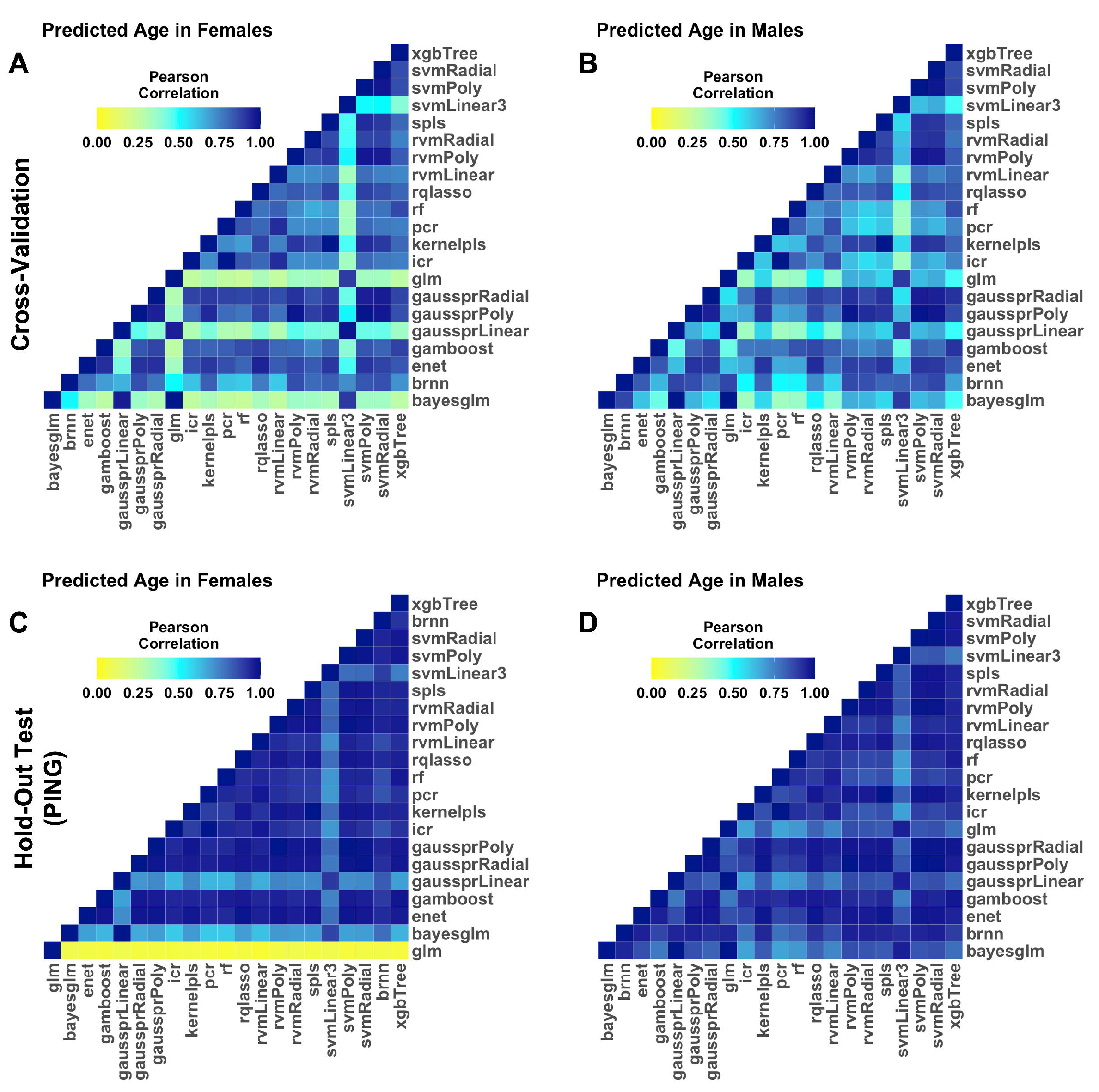
Pairwise Correlations of the Predicted Age of the 21 Algorithms. Figure demonstrates correlations between predicted age as estimated by different models in the females and males in the cross-validation set (Panels A and B), and in females and males in the hold-out Pediatric Imaging, Neurocognition, and Genetics Data Repository (PING) dataset (Panels C and D). The different algorithms are referenced by the function used for their implementation. Bayesian Generalized Linear Model (bayesglm); Bayesian Regularized Neural Network (brnn); Elastic Net Regression (enet);Generalized Additive Model with Boosting (gamboost);: Gaussian Processes Regression Linear (gaussprLinear); Gaussian Processes Regression Polynomial (gaussprPoly); Gaussian Processes Regression Radial (gaussprRadial); Generalized Linear Model (glm); Independent Component Regression (icr); Kernel Partial Least Squares (kernelpls); Principal Component Regression (pcr); Random Forest (rf); Quantile Regression with LASSO penalty (rqlasso); Relevance Vector Machine-Linear (rvmLinear3); Relevance Vector Machine-Polynomial (rvmPoly); rvmRadial: Relevance Vector Machine-Radial; Sparse Partial Least Squares (spls); Support Vector Regression-Linear (svmeLinear3); Support Vector Regression-Polynomial (svmPoly); Support Vector Regression-Radial (svmRadial); Extreme Gradient Boosting (xgbTree)

**Figure 3.**
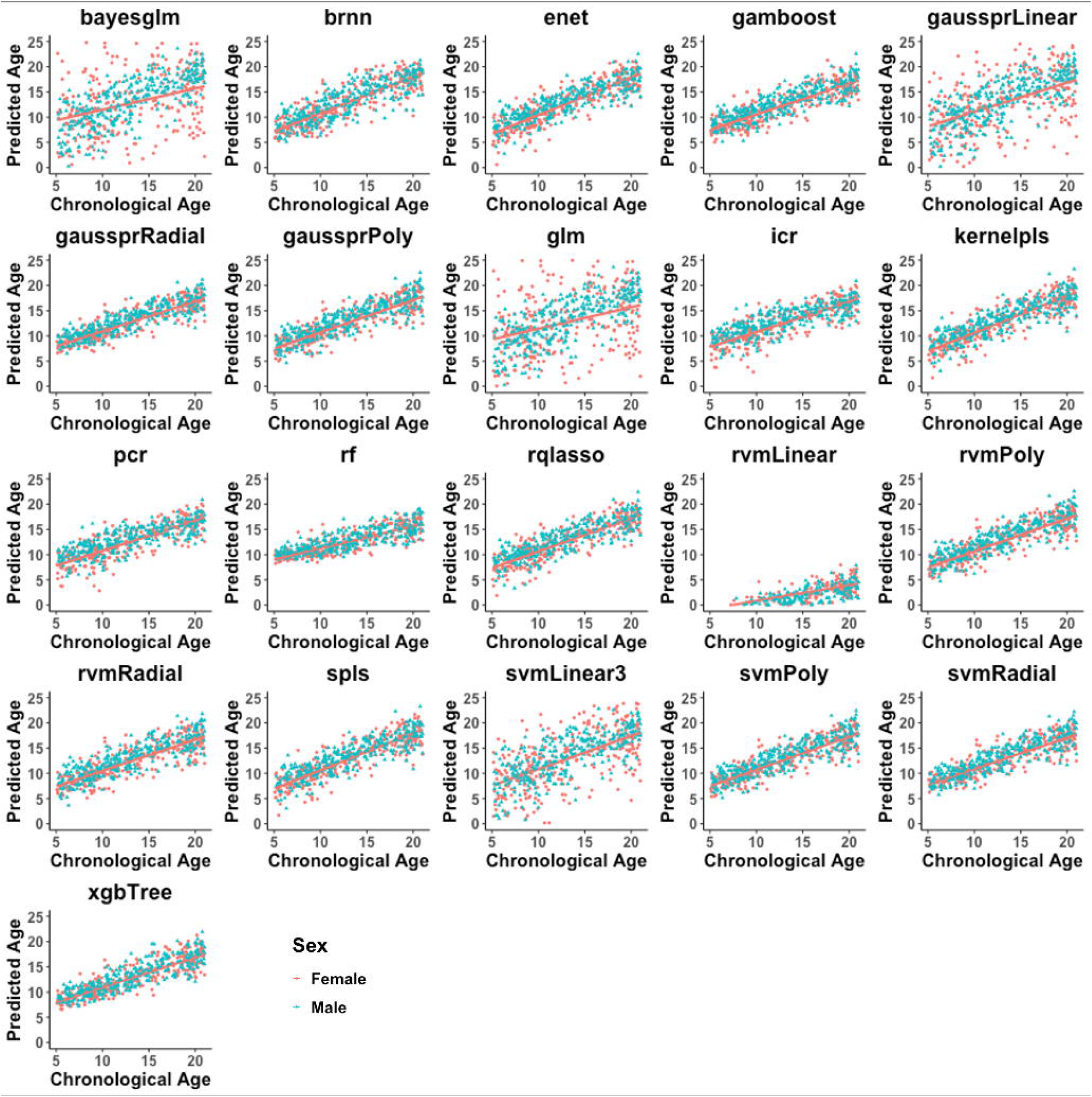
Correlations Between Chronological Age (Years) and Predicted Age Across 21 Algorithms in the PING Dataset. The figure shows the correlation of chronological age with sMRI-age in each of the 21 algorithms tested in males and females in the Pediatric Imaging, Neurocognition, and Genetics Data Repository (PING). The different algorithms are referenced by the function used for their implementation. Bayesian Generalized Linear Model (bayesglm); Bayesian Regularized Neural Network (brnn); Elastic Net Regression (enet);Generalized Additive Model with Boosting (gamboost);: Gaussian Processes Regression Linear (gaussprLinear); Gaussian Processes Regression Polynomial (gaussprPoly); Gaussian Processes Regression Radial (gaussprRadial); Generalized Linear Model (glm); Independent Component Regression (icr); Kernel Partial Least Squares (kernelpls); Principal Component Regression (pcr); Random Forest (rf); Quantile Regression with LASSO penalty (rqlasso); Relevance Vector Machine-Linear (rvmLinear3); Relevance Vector Machine-Polynomial (rvmPoly); rvmRadial: Relevance Vector Machine-Radial; Sparse Partial Least Squares (spls); Support Vector Regression-Linear (svmeLinear3); Support Vector Regression-Polynomial (svmPoly); Support Vector Regression-Radial (svmRadial); Extreme Gradient Boosting (xgbTree)

### 3.2 Computational speed and memory usage of each algorithm

The highest maximum memory usage was observed while training the Bayesian regularized neural networks, the boosted generalized additive model, and XGBoost algorithms (in that order). Highest average memory usage was seen with Bayesian regularized neural networks, the boosted generalized additive model, and the quantile regression with LASSO penalty. Bayesian regularized neural networks, XGBoost, and SVR with a polynomial kernel engaged CPU for the longest time. The generalized linear model, Kernel-partial least squares and principal component regression were the fastest algorithms with the lowest memory usage. Among the algorithms that performed best nonlinear kernelized versions had a favorable memory-computational speed profile (Supplemental Table 3).

### 3.3 BrainAGE and BrainAGE_corr_ for the three best performing algorithms

In Table 2 we report the BrainAGE and BrainAGE_corr_ derived from the three best performing algorithms. The corresponding values for all algorithms are shown in Supplemental Table 4. The values presented refer to the models’ performance in the ABCD and the PING samples using the optimized model parameters in the training phase. There was a significantly negative correlation between BrainAGE and chronological age but not for BrainAGE_corr_, as this association was mitigated by applying age-bias correction (Supplemental Table 4, Supplemental Figures 4-8).

**Table 2.** BrainAGE and BrainAGE_corr_ in the ABCD and PING Sample.

|  | ABCD |  |  |  |  | PING |  |  |
| --- | --- | --- | --- | --- | --- | --- | --- | --- |
| Algorithm (function name in caret package) | Males BrainAGE | Males BrainAGE<sub>corr</sub> | Females BrainAGE | Females BrainAGE<sub>corr</sub> | Males BrainAGE | Males BrainAGE<sub>corr</sub> | Females BrainAGE | Females BrainAGE<sub>corr</sub> |
| Extreme Gradient Boosting (xgbTree) | 1.33 (1.53) | 0.22 (1.50) | 0.73 (1.52) | -0.15 (1.49) | 0.06 (2.51) | 0.05 (1.66) | -0.28 (2.68) | -0.19 (1.80) |
| Random Forest Regression (rf) | 1.55 (1.46) | -0.03 (1.38) | 0.97 (1.34) | -0.37 (1.27) | -0.13 (3.10) | -0.19 (1.42) | -0.43 (3.10) | -0.34 (1.50) |
| Support Vector Regression -Radial (svmRadial) | 1.44 (1.57) | 0.56 (1.55) | 1.04 (1.57) | 0.21 (1.55) | 0.24 (2.40) | 0.26 (1.73) | -0.22 (2.62) | -0.16 (1.88) |

ABCD: Adolescent Brain Cognitive Development; MAE<sub>T</sub>: Mean Absolute Error; PING: Pediatric Imaging, Neurocognition, and Genetics Data Repository; Values shown as mean (standard deviation); BrainAGE<sub>corr</sub>: Age-bias corrected BrainAGE

### 3.4 Feature importance in brain-age prediction in the three best performing algorithms

Feature importance, as inferred by their SVs, varied considerably across models specified with XGBoost, RF regression, and SVR with the RBF kernel (Figure 4, Supplemental Table 5). The values presented refer to the models’ performance in the training sample using the optimized model parameters. In RF regression, a few regions made very large contributions, with minimal contributions from other regions. In SVR with the RBF kernel, most features contributed to the model although the contribution of each feature was small. The profile of feature contributions in XGBoost was intermediate between the other two algorithms.

**Figure 4.**
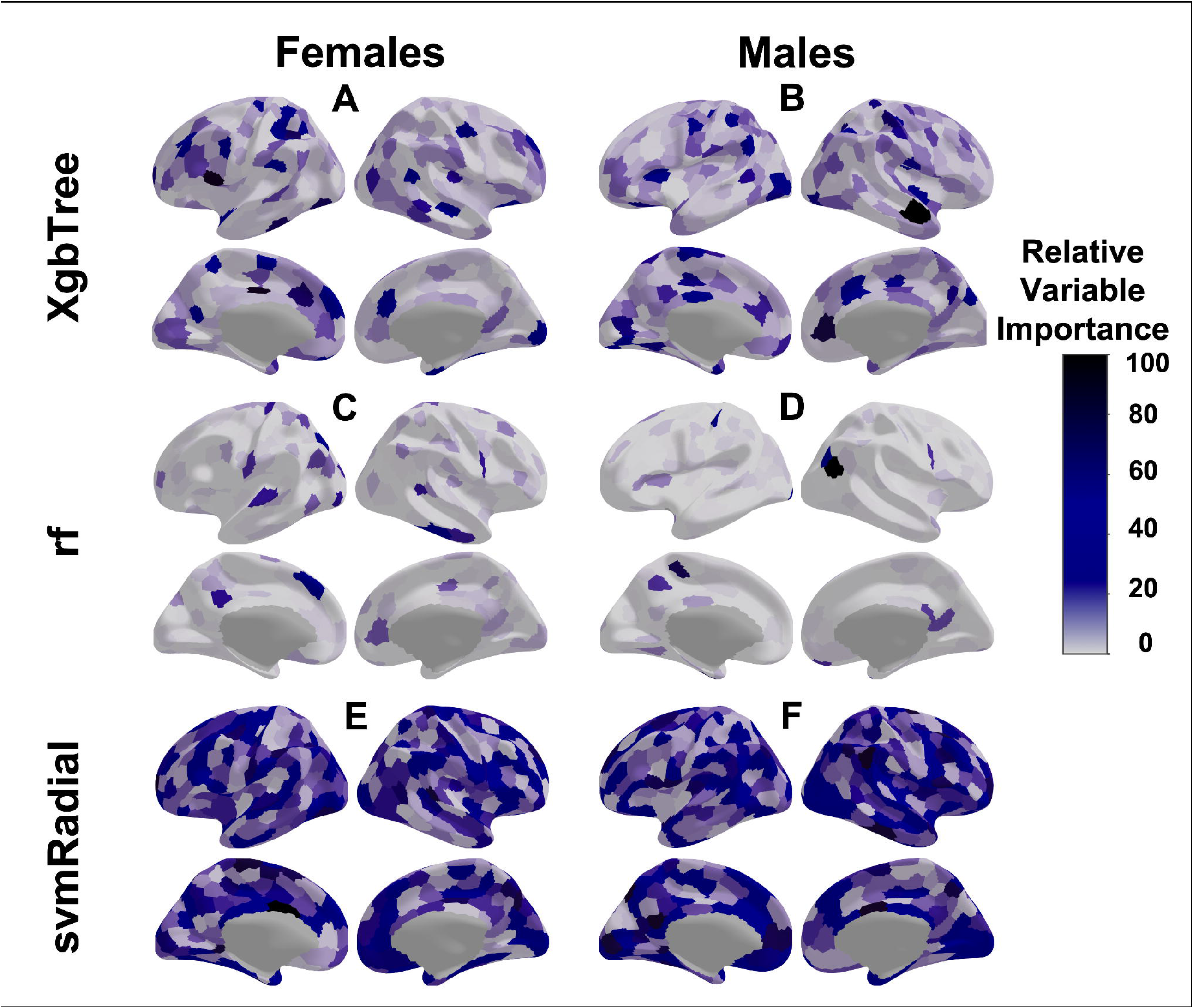
Relative Importance of Neuroimaging Features for Age Prediction in the Three Best Performing Algorithms. The figure shows the relative variable importance of the features of the 400-parcel Schaefer Atlas for age-prediction based on their Shapley Values in females (Panel A) and males (Panel B). The relative importance values shown were rescaled such that the feature with the maximum average absolute Shapley Value in each model was assigned a value of 100. The algorithms are referenced by the function used for their implementation: Random Forest Regression (rf); Support Vector Regression-Radial (svmRadial); Extreme Gradient Boosting (xgbTree)

### 3.5 Sensitivity and supplemental analyses for the three best performing algorithms

Sex: Application of parameters from models trained on males to the entire sample, yielded marginally higher BrainAGE values for females than males (maximum difference across models=0.2 years) (Supplemental Tables 6 and 7) Similarly, application of parameters from models trained on females to the entire sample yielded higher BrainAGE for females than males (maximum difference across models=0.6 years) (Supplemental Tables 6 and 7).

Parcellation scheme and number of input features: The different parcellation schemes had minimal influence on the MAE in any of the three best performing algorithms. The number of features in the range examined (136-2,000) had minimal impact on MAE and notably the Schaefer-1000 parcellation did not outperform the Schaefer-400 parcellation used in the main analyses (Figure 5A). The same pattern was seen in females and males in the ABCD and PING datasets (Supplemental Figures 9 and 10).

**Figure 5.**
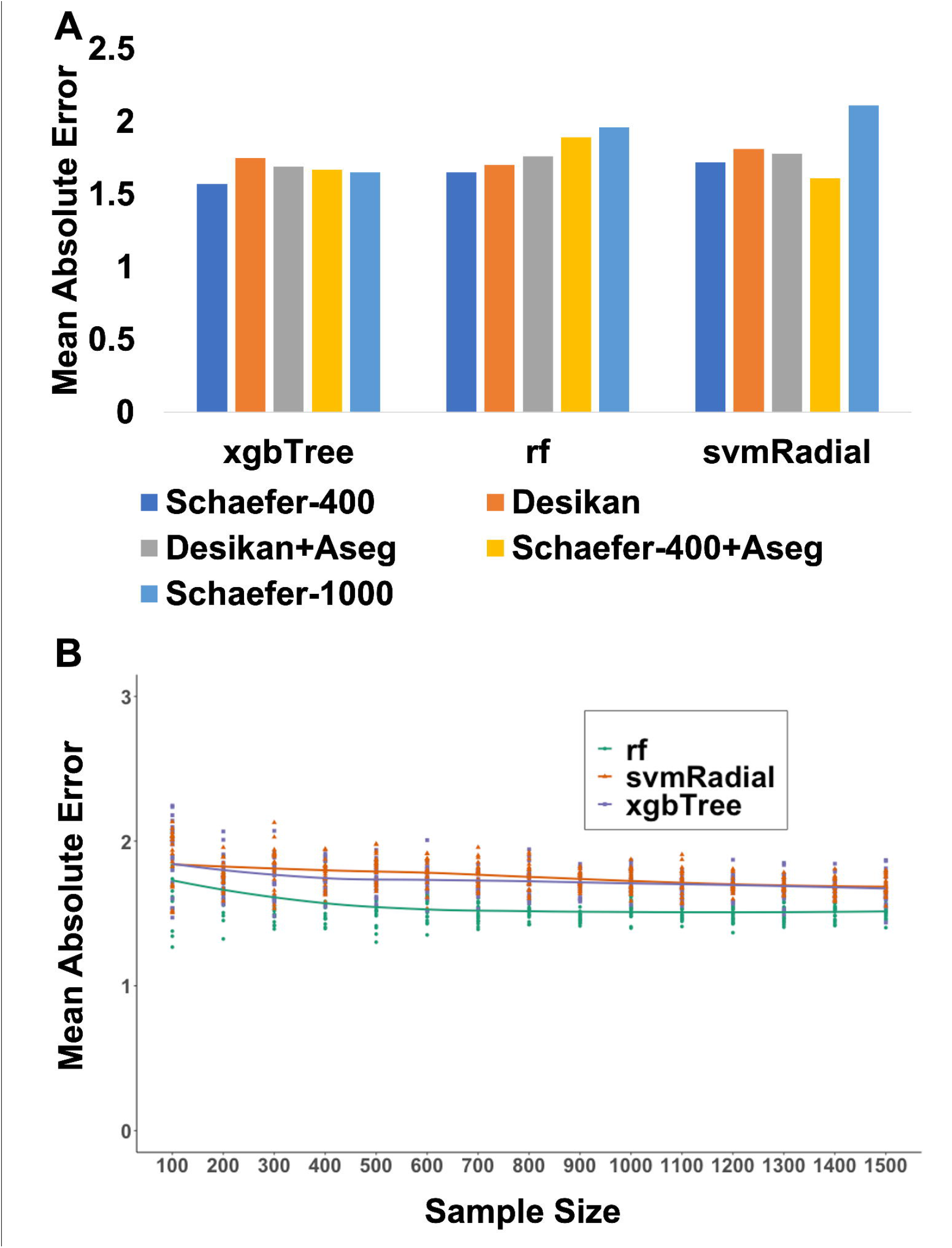
Mean Absolute Error (MAE) as a Function of Parcellation Scheme and Sample Size. **Panel A**: The mean absolute error for the three best performing algorithms as a function of parcellation scheme in male participants from the Adolescent Brain Cognitive Development (ABCD) study. The corresponding information from female ABCD participants and the PING sample are shown in Supplemental Figures 8 and 9); **Panel B:** Mean absolute error of each of the three best performing algorithms as a function of sample size in the ABCD sample; model parameters for each algorithm were obtained by randomly resampling the training dataset without replacement generating subsamples of 100 to 1500. The algorithms are referenced by the function used for their implementation: Random Forest Regression (rf); Support Vector Regression-Radial (svmRadial); Extreme Gradient Boosting (xgbTree)

Sample size: MAE_T_ improved in line with sample size increase up to a size of 500 participants and it plateaued thereafter. The corrected MAE_T,_ on the other hand sowed limited change across sample size (Figure 5B, Supplemental Figure 11).

Number of cross-validation folds and repeats: In the main analyses, we used 5-fold repeats and 5-fold cross-validations. Using 10 instead of 5 folds and repeats did not improve performance and in the case of XGBoost, we noted markedly worse performance in the MAE_T_ (Supplemental Table 8).

Effect of extreme outliers in the test set: In the PING dataset, spearman’s correlation coefficients between the number of outliers and the MAE derived from the three best performing algorithms was small (all rho<0.15). In the ABCD dataset, the corresponding values were of similar magnitude with a maximum rho of 0.25 for RF regression. The magnitude of these associations was reduced when using age-bias-corrected MAE (max rho<0.2). Among the three best performing models, the performance of SVR with the RBF kernel was the least impacted by extreme outliers (Supplemental Table 9).

## 4. Discussion

In the present study, we undertook a comprehensive comparison of machine learning algorithms for sMRI-based age prediction as a proxy for the biological age of the brain in youth. We identified three algorithms, namely XGBoost, RF regression, and SVR with the RBF kernel, that outperformed all others in terms of accuracy while being computationally efficient. Notably we also show that sMRI-based age prediction was suboptimal in models using linear algorithms.

Linear algorithms consistently underperformed compared to other algorithms probably because of the multicollinearity of the neuroimaging data, as suggested by the relative better performance of those linear algorithms that are based on covariance (such as SPLS regression or PCA regression). Further, the general underperformance of linear models may also reflect the fact that they do not account for nonlinear and interactive associations between brain imaging features and age.

As predicted XGBoost and RF regression, which are both ensemble tree-based algorithms, performed well in terms of their accuracy and generalizability to unseen samples. A decision tree is a machine learning algorithm that partitions the data into subsets based on conditional statements. Although each tree has generally low predictive performance, their combination (ensemble) improves generalizability without sacrificing accuracy (Qi, 2012). Further advantages of these methods, particularly in the context of neuroimaging datasets, is that they are nonparametric, they do not involve assumptions about the distribution for the data and can account for nonlinear effects and interactions, which may be particularly relevant in modeling developmental brain-age. They also require minimal preparation of the input sMRI features as they can handle multicollinear data without losing accuracy. Notably, these algorithms were relatively insensitive to the number of the neuroimaging features, when the size of the feature sets ranged between 136 to 2000, with the mid-point feature set (n=841) being relatively better.

Similar considerations applied to SVR with RBF algorithm which had the additional advantage of being particularly robust to outliers. This may represent a particular advantage of this algorithm for studies with relatively small dataset where strict rules for outlier exclusion may result in significant data loss. Despite similar performance in terms of accuracy, the sMRI features contributing to age prediction differed across the three best-performing algorithms. Relative to the other two algorithms, more features contributed to age prediction in SVR with RBF which may contribute to its robustness to outliers.

A major concern in neuroimaging research is the effect of site on the generalizability of ML models (Dockes et al., 2021; Solanes et al., 2021). Sites may differ in terms of scanner infrastructure, acquisition protocols and neuroimaging feature extraction pipelines as well as sample composition. Here the post-acquisition extraction of neuroimaging features was undertaken for all cohorts using the same pipeline which may have reduced variability in the neuroimaging feature set. However, all other parameters differed between cohorts (and between recruitment sites within cohorts). Yet the three best performing algorithms showed excellent generalizability to the hold-out datasets, which is likely to reflect the robustness of these algorithms. Additionally, the inclusion of observations from multiple sites in the training dataset may have forced the ML algorithms to select and weight features that are robust to site differences, therefore reducing the dependence of the model on the effects of site.

We observed a lower MAE for females compared to males across most models. This has been reported in prior studies (Brouwer et al., 2021; Wierenga et al., 2019; Wierenga et al., 2022), and can be attributed to either biological differences, i.e., female brain showing less variability or confounding, i.e., males may move more, on average, than females which could make their brain measurements less accurate. Brouwer and colleagues (Brouwer et al., 2021) demonstrated that in individuals ages 9-23, females have higher sMRI-derived BrainAGE than their male counterparts. The same pattern was reported by Tu and colleagues (Tu et al., 2019) using sMRI data from 118 males and 147 females, aged 5-18, from the NIH MRI Study of Normal Brain Development. In these studies, as well as in ours, sex differences in BrainAGE are small and within the range of the MAE for brain-predicted age.

We acknowledge that the list of algorithms evaluated is not exhaustive but provides a good coverage of the many models that are currently available. We were unable to account for potential influences of race and ethnicity as such information was either absent or not uniformly coded in the cohorts used for model training. Based on the racial constitution of the general population, in the countries of the recruitment sites, we anticipate an over-representation of white individuals. As more data becomes available on other racial/ethnic groups, it should be possible to address this issue in future studies.

In summary, using a wide range of ML algorithms on geographically diverse datasets of young people, we showed that tree-based followed by nonlinear kernel-based algorithms offer robust, accurate, and generalizable solutions for predicting age based on brain morphological features. Findings of the present study can be used as a guide for quantifying brain maturation during development and its contribution to functional and behavioral outcomes.

## Acknowledgements

Data used in the preparation of this article were obtained from the Adolescent Brain Cognitive Development (ABCD) Study (https://abcdstudy.org), held in the NIMH Data Archive (NDA). This is a multisite, longitudinal study designed to recruit more than 10,000 children age 9-10 and follow them over 10 years into early adulthood. The ABCD Study is supported by the National Institutes of Health and additional federal partners under award numbers U01DA041022, U01DA041028, U01DA041048, U01DA041089, U01DA041106, U01DA041117, U01DA041120, U01DA041134, U01DA041148, U01DA041156, U01DA041174, U24DA041123, U24DA041147, U01DA041093, and U01DA041025. A full list of supporters is available at https://abcdstudy.org/federal-partners.html. A listing of participating sites and a complete listing of the study investigators can be found at https://abcdstudy.org/scientists/workgroups/. ABCD consortium investigators designed and implemented the study and/or provided data but did not necessarily participate in analysis or writing of this report. This manuscript reflects the views of the authors and may not reflect the opinions or views of the NIH or ABCD consortium investigators. The ABCD data repository grows and changes over time. Dr. A.M. received support from the National Institute of Mental Health T32-MH122394. Dr S.F. received support from the National Institute of Mental Health (R01-MH113619). This work was supported in part through the computational resources and staff expertise provided by Scientific Computing at the Icahn School of Medicine at Mount Sinai.

## Author statement

AM: Conceptualization, Methodology, Investigation, Data Curation, Formal analysis, Writing - Original Draft, Writing - Review & Editing

SF: Supervision, Conceptualization, Writing - Original Draft, Writing - Review & Editing

RSK: Supervision, Conceptualization, Writing - Review & Editing

HCW: Supervision, Conceptualization, Review & Editing

DCG: Supervision - Review & Editing

PMT: Supervision - Review & Editing

