## Supplemental file for "Systematic Evaluation of Machine Learning Algorithms for Neuroanatomically-Based Age Prediction in Youth"

**Supplementary Material**

**Supplemental Methods**

**Samples**

Description of the samples

Details of Excluded Participants per Sample

Supplemental Table 1. Characteristics of Included Datasets

Supplemental Figure 1. Age and Sex Distribution of the Training Dataset

Supplemental Figure 2. The Schaefer 400 parcellation scheme

**Supplemental Results**

Supplemental Table 2. Algorithm Accuracy Across Cross-Validations in the Training Datasets

Supplemental Table 3. Computational Costs of the Algorithms In the Training Dataset

Supplemental Table 4. BrainAGE and BrainAGE_corr_ in the Hold-Out Datasets

Supplemental Figure 3. Correlations Between Predicted and Chronological Age (years) in the Cross-Validation Sets.

Supplemental Figure 4. Correlations Between Brain Age Gap Estimation (BrainAGE) and Chronological Age (years) in the Cross-Validation Sets

Supplemental Figure 5. Correlations Between Age-Bias Corrected Brain Age Gap Estimation (BrainAge_corr_) and Chronological Age (years) in the Cross-Validation Sets.

Supplemental Figure 6. Correlations Between Brain Age Gap Estimation (BrainAGE) and Chronological Age (years) in the PING dataset.

Supplemental Figure 7. Correlations Between Age-Bias Corrected Brain Age Gap Estimation (BrainAGE_corr_) and Chronological Age (years) in the PING Dataset.

Supplemental Figure 8. Age Bias-Corrected Mean Absolute Error (MAE_corr_) of the 21 Algorithms.

Supplemental Table 5. Top 20 Features Contributing to Age Prediction in the Three Best-Performing Algorithms by Sex

Supplemental Table 6. Mean absolute error (MAE) and Age-Bias Corrected Mean Absolute Error (MAE_corr_) of Models Trained in One Sex and Tested on the Other

Supplemental Table 7. BrainAGE and BrainAGE_corr_ Derived from Models Trained on One Sex and Tested in the Other

Supplemental Figure 9. ABCD Dataset: Mean Absolute Error per Parcellation Scheme in Each of the Three Best Performing Algorithms

Supplemental Figure 10. PING Dataset: Mean Absolute Error per Parcellation Scheme in Each of the Three Best Performing Algorithms

Supplemental Figure 11. Age-Bias Corrected Mean Absolute Error (MAE_corr_) of the Three Best Performing Algorithms as a Function of Sample Size

Supplemental Table 8. Mean Absolute Error (MAE) and Age-Bias corrected Mean Absolute Error (MAE_corr_) Between 5 and 10 Repeats of the Tenfold Cross-Validation in the Three Best Performing Algorithms

Supplemental Table 9. Spearman's Correlation Coefficients Between the Number of Outliers, Mean Absolute Error (MAE) and Age-Bias Corrected Mean Absolute Error (MAE_corr_) for the Three Best Performing Algorithms

**Supplemental References**

**Supplemental Methods**

**Samples**

***Autism Brain Imaging Data Exchange (ABIDE) and ABIDE II***

ABIDE is a multi-site consortium aggregating and openly sharing 1112 multimodal neuroimaging data and phenotypic information from 539 individuals with Autism Spectrum Disorder (ASD) and 573 age-matched healthy comparison individuals (age range: 7–64 years) (<http://fcon_1000.projects.nitrc.org/indi/abide/>).

ABIDE II is an extension of the prior study and has released neuroimaging and phenotypic data from 521 individuals with ASD and 593 comparison subjects (age range: 5-64 years) (http://fcon_1000.projects.nitrc.org/indi/abide/abide_II.html). Details of these initiatives have been published by (Di Martino et al., 2017; Di Martino et al., 2014).

***Attention Deficit Hyperactivity Disorder (ADHD)-200***

The ADHD-200 Consortium (http://fcon_1000.projects.nitrc.org/indi/adhd200/#) is a multi-site consortium aggregating and sharing multimodal neuroimaging data and phenotypic information from 261 individuals diagnosed with Attention Deficit Hyperactivity Disorder (ADHD) (aged 7-21 years) and 430 age-matched healthy comparisons individuals. Details of the Consortium have been published by the ADHD consortium (ADHD-200 Consortium, 2012).

***Child Mind Institute-Healthy Brain Network (CMI-HBN)***

The Healthy Brain Network (HBN) is an ongoing project by the Child Mind Institute (CMI) aiming to collect and share neuroimaging, blood-based and phenotypic data from 10,000 participants (ages 5–21) (<http://fcon_1000.projects.nitrc.org/indi/cmi_healthy_brain_network/>) living in New York and surrounding areas. To-date, the CMI-HBN has released data of 2168 individuals. Details of the CMI-HBN have been by published by (Alexander et al., 2017)

***Human Connectome Project- Development (HCP-D)***

The Human Connectome Project-Development (HCP-D) is an ongoing study collecting and sharing multimodal neuroimaging data and phenotypic information from individuals aged 5 to 21 years (https://www.humanconnectome.org/study/hcp-lifespan-development). To-date data on 652 participants have been released. Details of the HCP-D have been published by Harms et al., 2018 (Harms et al., 2018).

***Adolescent Brain Cognitive Development (ABCD)*** ***study***

The Adolescent Brain Cognitive Development (ABCD) study (https://nda.nih.gov/abcd/) is a prospective multisite study that aims to collect and share multimodal imaging and associated phenotypes and genotypes from a population–based sample of 11,875 pre-adolescents (age-range: 9 to10 years) to be followed-up prospectively for up to 10 years. The baseline data release provides information on ABCD participants. A subset of the baseline data was used as the held-out-test set in this study. Details of the ABCD study design and of the neuroimaging acquisition and quality assurance procedures have been published by (Casey et al., 2018; Hagler et al., 2019)

***Pediatric Imaging, Neurocognition, and Genetics (PING) Study***

The Pediatric Imaging, Neurocognition, and Genetics (PING) study is a cross-sectional study of pf 1,493 typically developing children aged 3-21 (http://pingstudy.ucsd.edu/). The data involves clinical measures, and brain MRI and genetic data. A subset of this sample (N=754) had available raw T1-weighted images was used as the held-out-test set in this study. Details of the PING study design and of the neuroimaging acquisition and quality assurance procedures have been published by (Jernigan et al., 2016).

**Details of Excluded Participants per Sample in the Training Set**

***Data selection for the training dataset***: Participant selection from the ABIDE, ABIDE II, ADHD-200, HCP-D and CMI-HBN followed a two-step process. First, in each sample participants were excluded if they had (a) a psychiatric or neurological disorder; and (b) they had missing values in their parcellation. Second, after collating the data from all participants selected at the first step, we further excluded participants if more than 5% of their parcellation features had extreme values defined as being higher or lower than 3 median absolute deviations from the median of the entire training sample. Based on these criteria, the following participants were excluded:

**ABIDE**: 539 participants (48% of the total ABIDE sample) were excluded because of psychiatric morbidity; 100 (9% of the total ABIDE sample) participants were excluded due to the absence of T1-weighted images; 4 (0.3% of the total ABIDE sample) were excluded because of outlier values.

**ABIDE II:** 521 participants (47% of the total ABIDE II sample) were excluded because of psychiatric morbidity; 143 (13% of the total ABIDE sample) participants were excluded due to the absence of T1-weighted images; 17 (1% of the total ABIDE II sample) participants were excluded because of failed parcellation of their T1-weighted images; 5 (0.4% of the total ABIDE II sample) were excluded because of outlier values.

**ADHD-200:** 261 participants (38% of the total ADHD-200 sample) were excluded because of psychiatric morbidity; 40 (6% of the total ABIDE sample) participants were excluded due to the absence of T1-weighted images; 4 (0.6% of the total ADHD-200 sample) were excluded because of outlier values.

**CMI-HBN:** 1876 participants (86% of the total CMI-HBN sample) were excluded because of psychiatric morbidity; 73 (3 % of the total CMI-HBN sample) participants were excluded due to the absence of T1-weighted images; 14 (0.6% of the total CMI-HBN sample) were excluded because of outlier values.

**HCP-D**: None of the HCP-D participants were excluded

***Data selection for the held-out independent dataset:*** We followed the same process as for the training set.

**ABCD**: 7511 participants (63% of the total ABCD sample) were excluded because of psychiatric morbidity; 286 (2% of the total ABCD sample) participants were excluded because of failed parcellation of their T1-weighted images. We didn’t perform outlier exclusion on the ABCD dataset.

**PING**: of 1,493 participants, 739 (49% of the total PING sample) were excluded because they had no available raw T1-weighted images. 105 (7% of the total PING sample) participants were excluded because of a learning disability or psychopathology. Two participants (0.1% of the total PING sample) were excluded because their T1-weighted images failed to process. 54 (4%) participants were further excluded because they were younger than 5 years old.

| **Supplemental Table 1. Characteristics of Included Cohorts** | | | | |
| --- | --- | --- | --- | --- |
| **Cohort** | **Sample Size** | **Females N (%)** | **Mean Age (Range) (Years)** | **Scanner Vendor and Magnet Strength** |
| **Datasets Used In the Training Set** | | | | |
| **ABIDE** | 436 | 83 (19%) | 13.6 (6.5-21) | GE_MR750_3T, GE_Signa_3T; Philips_Achieva_3T; Philips_Interna_3T; Siemens_Allegra_3T; Siemens_TrioTim_3T, Siemens_Veiro_3T |
| **ABIDE-II** | 427 | 139 (32.5%) | 11.5 (5.9-21) | GE_MR750_3T; Philips_Achieva_1.5T; Philips_Achieva_3T; Philips_Ingenia_3T, Siemens_Allegra_3T, Siemens_Skyra_3T; Siemens_TrioTim_3T |
| **ADHD-200** | 385 | 206 (53.5%) | 12.1 (7.1-20.5) | Philips_Achieva_3T, Siemens_Allegra_3T, Siemens_Avanto_1.5T, Siemens_TrioTim_3T |
| **CMI-HBN** | 205 | 90 (43.9%) | 10.0 (5-21 | Siemens_Avanto_1.5T; Siemens_Prisma_3T; Siemens_TrioTim_3T |
| **HCP-D** | 652 | 351 (53.8%) | 14.4 (5.6-22) | Siemens_Prisma_3T |
| **Held-out test datasets** | | | | |
| **ABCD** | 4078 | 2126 (52%) | 9.9 (0.6) | Siemens_Prisma_3T, Siemens_VE11B-C_3T; Philips_Achieva_3T; Philips_dStream_3T; Philips_Ingenia_3T  GE_MR750_3T; GE_DV25-26_3T |
| **PING** | 594 | 295 (49.6%) | 12.9 (4.8) | Philips_Achieva_3T  GE_SIGNA_3T  Siemens_TrioTim_3T  Siemens_TrioTim_3T  GE_Discovery_3T  MR750_3T |

**Supplemental Figure 1. Sex And Age Distribution in the Training Dataset.**

The sex and age distribution of participants the collated cohorts that comprise the training set (ABIDE, ABIDE II, ADHD-200, HCP-D; CMI; N=2,105)

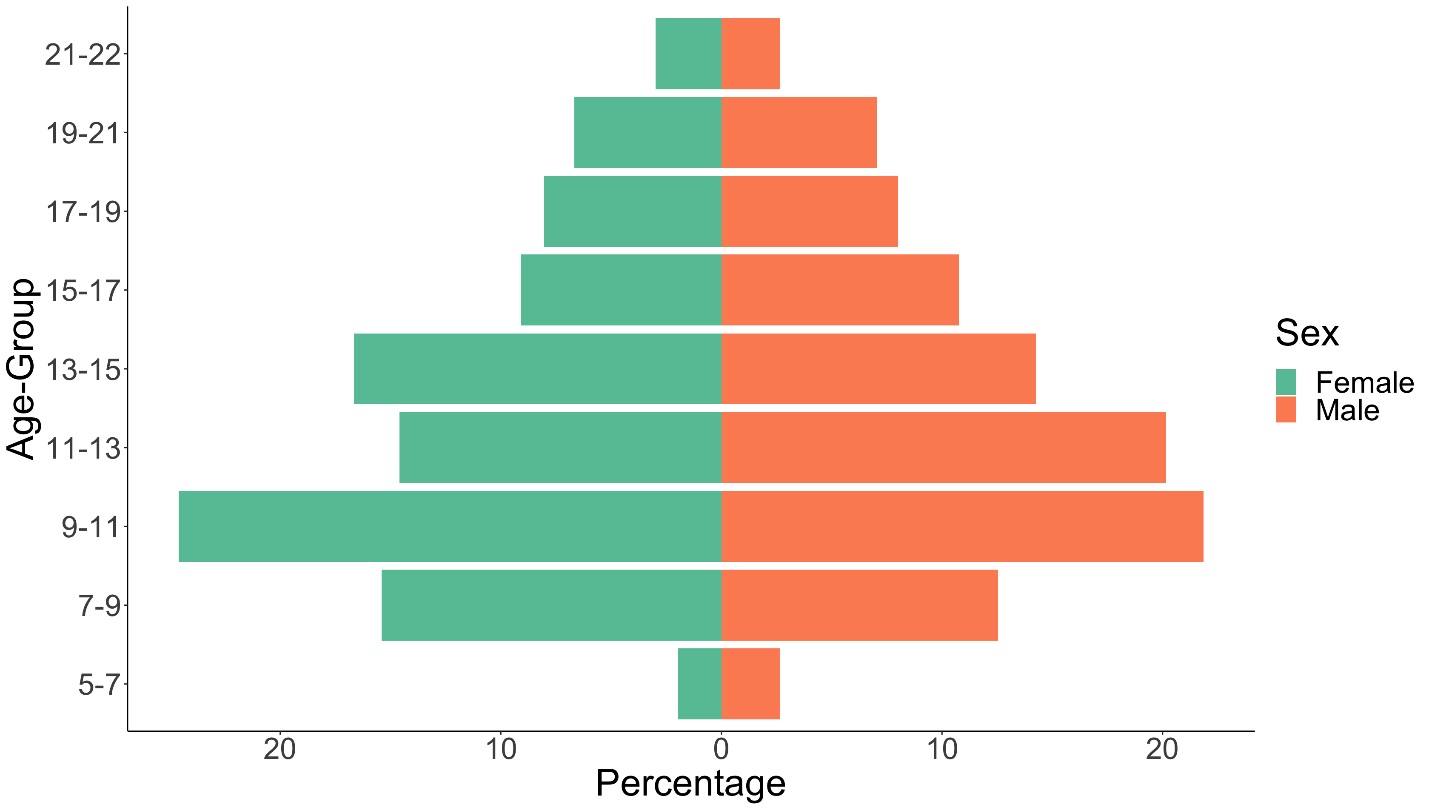

**Supplemental Figure 2. The Schaefer 400 parcellation scheme**
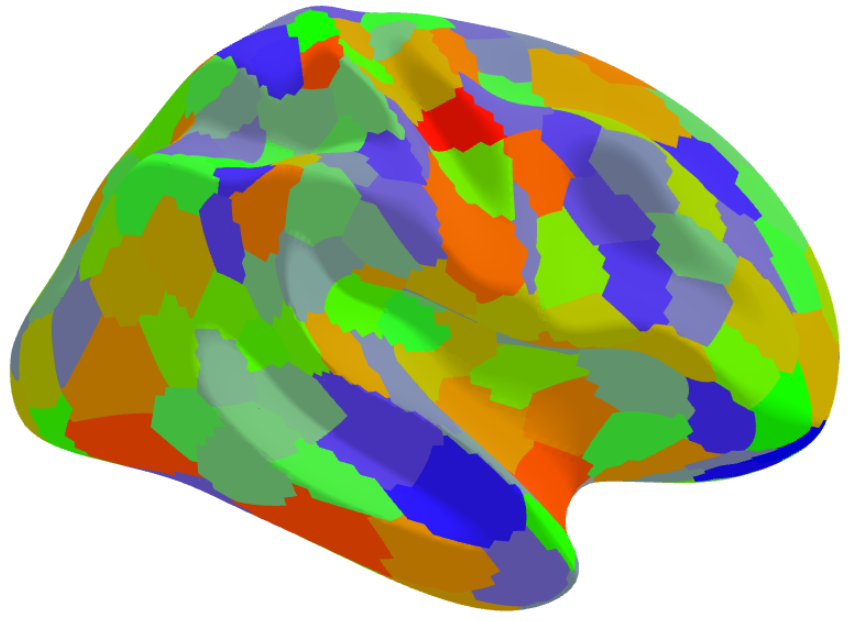

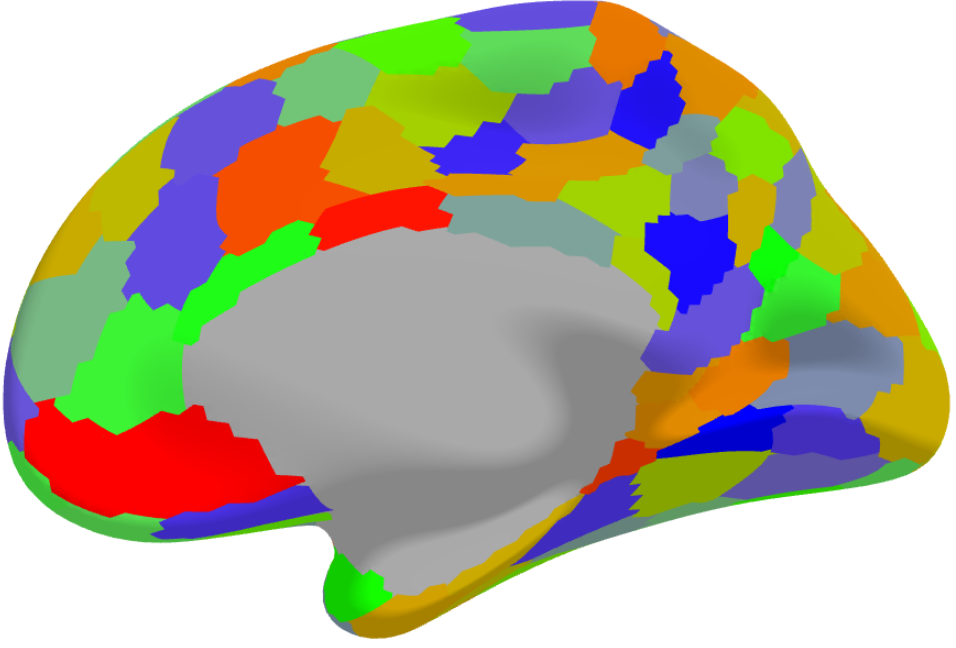

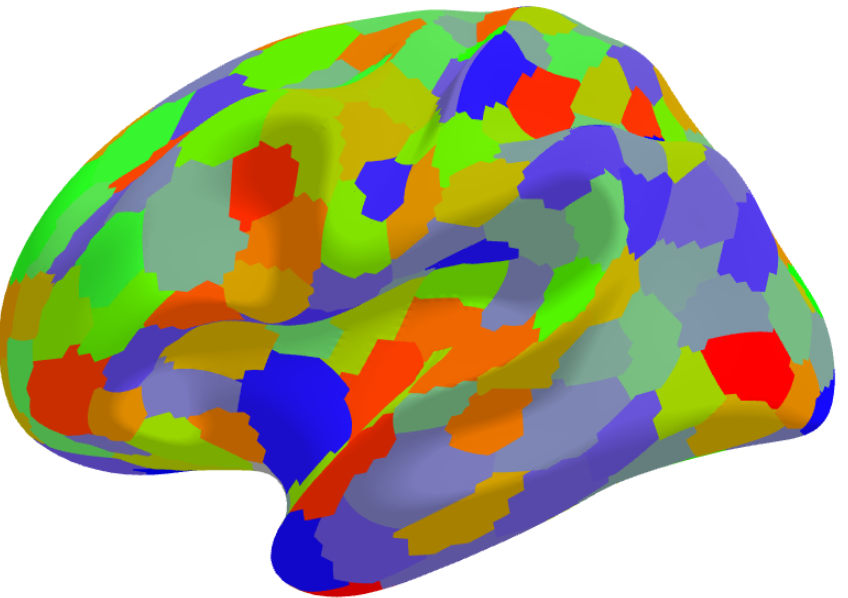

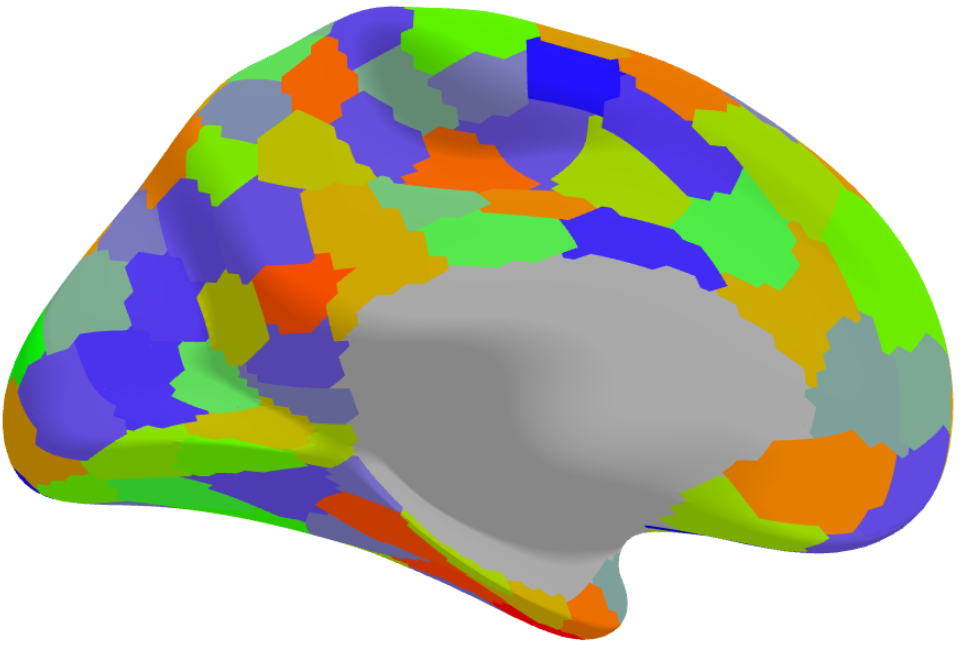

**Supplemental Results**

| **Supplemental Table 2. Algorithm Accuracy Across Cross-Validations in the Training Datasets** | | | |
| --- | --- | --- | --- |
| **Algorithm (function name in caret package)** | **MAE_cv_** | **RMSE_cv_** | **Correlation_cv_** |
| **Males** | | | |
| **Extreme Gradient Boosting (xgbTree)** | 1.7 | 2.18 | 0.83 |
| **Random Forest Regression (rf)** | 1.99 | 2.52 | 0.77 |
| **Support Vector Regression -Radial (svmRadial)** | 1.6 | 2.1 | 0.84 |
| **Support Vector Regression -Polynomial (svmPoly)** | 1.63 | 2.14 | 0.83 |
| **Relevance Vector Regression-Polynomial (rvmPoly)** | 1.65 | 2.16 | 0.83 |
| **Gaussian Processes Polynomial (gaussprPoly)** | 1.63 | 2.13 | 0.83 |
| **Gaussian Processes Radial (gaussprRadial)** | 1.71 | 2.18 | 0.83 |
| **Generalized Additive Model with Boosting (gamboost)** | 1.81 | 2.31 | 0.80 |
| **Sparse Partial Least Squares (spls)** | 1.77. | 2.26 | 0.81 |
| **Kernel Partial Least Squares (kernelpls)** | 1.77 | 2.26 | 0.81 |
| **Elastic Net Regression (enet)** | 1.73 | 2.21 | 0.82 |
| **Quantile Regression with LASSO penalty (rqlasso)** | 1.77 | 2.26 | 0.81 |
| **Relevance Vector Regression-Radial (rvmRadial)** | 1.71 | 2.23 | 0.82 |
| **Bayesian Regularized Neural Network (brnn)** | 1.93 | 2.44 | 0.79 |
| **Independent Component Regression (icr)** | 2.19 | 2.72 | 0.71 |
| **Principal Component Regression (pcr)** | 2.19 | 2.72 | 0.71 |
| **Support Vector Regression-Linear (svmLinear3)** | 2.87 | 3.67 | 0.65 |
| **Gaussian Processes-Linear (gaussprLinear)** | 3.21 | 4.1 | 0.6 |
| **Generalized Linear Model (glm)** | 3.43 | 4.39 | 0.57 |
| **Bayesian Generalized Linear Model (bayesglm)** | 3.43 | 4.39 | 0.57 |
| **Relevance Vector**  **Machine-Linear (rvmRaidal)** | 12.89 | 13.14 | 0.76 |
| **Females** | | | |
| **Random Forest Regression (rf)** | 1.82 | 2.33 | 0.82 |
| **Extreme Gradient Boosting (xgbTree)** | 1.59 | 2.08 | 0.85 |
| **Support Vector Regression -Radial (svmRadial)** | 1.6 | 2.08 | 0.85 |
| **Support Vector Regression -Polynomial (svmPoly)** | 1.58 | 2.06 | 0.85 |
| **Gaussian Processes Polynomial (gaussprPoly)** | 1.61 | 2.09 | 0.85 |
| **Generalized Additive Model with Boosting (gamboost)** | 1.71 | 2.19 | 0.83 |
| **Relevance Vector Regression -Polynomial (rvmPoly)** | 1.63 | 2.13 | 0.84 |
| **Gaussian Processes Radial (gaussprRadial)** | 1.65 | 2.13 | 0.85 |
| **Relevance Vector Regression -Radial (rvmRadial)** | 1.75 | 2.22 | 0.83 |
| **Quantile Regression with LASSO penalty (rqlasso)** | 1.72 | 2.24 | 0.82 |
| **Independent Component Regression (icr)** | 1.93 | 2.42 | 0.79 |
| **Principal Component Regression (pcr)** | 1.93 | 2.43 | 0.79 |
| **Elastic Net Regression (enet)** | 1.7 | 2.18 | 0.84 |
| **Kernel Partial Least Squares (kernelpls)** | 1.72 | 2.19 | 0.83 |
| **Sparse Partial Least Squares (spls)** | 1.71 | 2.18 | 0.83 |
| **Bayesian Regularized Neural Network (brnn)** | 1.95 | 2.44 | 0.79 |
| **Support Vector Regression-Linear (svmLinear3)** | 3.3 | 4.18 | 0.59 |
| **Gaussian Processes-Linear (gaussprLinear)** | 4 | 5.08 | 0.51 |
| **Bayesian Generalized Linear Model (bayesglm)** | 4.6 | 5.84 | 0.46 |
| **Generalized Linear Model (glm)** | 123.87 | 160.56 | 0.1 |
| **Relevance Vector Machine-Linear (rvmLinear)** | 12.71 | 12.95 | 0.78 |
| LASSO: least absolute shrinkage and selection operator**;** MAE: Mean Absolute Error; RMSE: Root Mean Squared Error | | | |

| **Supplemental Table 3.** **Computational Costs of the Algorithms In the Training Dataset** | | | |
| --- | --- | --- | --- |
| **Algorithm** | **Max Memory (MB)** | **Average Memory (MB)** | **Run Time (Sec)** |
| **Bayesian Generalized Linear Model** | 16501 | 13854 | 991 |
| **Bayesian Regularized Neural Network** | 33835 | 29096 | 77099 |
| **Elastic Net Regression** | 18058 | 16240 | 5018 |
| **Generalized Additive Model with Boosting** | 48553 | 34016 | 5177 |
| **Gaussian Processes Regression Linear** | 14240 | 8138 | 175 |
| **Gaussian Processes Regression Polynomial** | 19908 | 16897 | 685 |
| **Gaussian Processes Regression Radial** | 18826 | 13187 | 490 |
| **Generalized Linear Model** | 11265 | 6892 | 240 |
| **Independent Component Regression** | 19556 | 12499 | 561 |
| **Kernel Partial Least Squares** | 12050 | 7060 | 135 |
| **Principal Component Regression** | 12631 | 7398 | 174 |
| **Random Forest** | 15333 | 14577 | 13582 |
| **Quantile Regression with LASSO penalty** | 19829 | 18472 | 10115 |
| **Relevance Vector Machine-Linear** | 14792 | 8869 | 173 |
| **Relevance Vector Machine-Polynomial** | 20450 | 18771 | 1656 |
| **Relevance Vector Machine-Radial** | 19981 | 16905 | 995 |
| **Sparse Partial Least Squares** | 16251 | 13907 | 948 |
| **Support Vector Regression-Linear** | 14455 | 13379 | 2343 |
| **Support Vector Regression -Polynomial** | 18352 | 16486 | 2385 |
| **Support Vector Regression -Radial** | 17645 | 16905 | 31811 |
| **Extreme Gradient Boosting** | 22238 | 18898 | 75015 |
| LASSO: least absolute shrinkage and selection operator | | | |

| **Supplemental Table 4.** **BrainAGE and BrainAGE_corr_ in the Hold-Out Datasets** | | | | |
| --- | --- | --- | --- | --- |
|  | **ABCD** | | **PING** | |
| **Algorithm (function name in caret package)** | **BrainAGE** | **BrainAGE_corr_** | **BrainAGE** | **BrainAGE_corr_** |
| **Males** | | | | |
| **Bayesian Generalized Linear Model** | 1.72 | 1.1 | 0.14 | 0.19 |
| **Bayesian Regularized Neural Network** | 1.47 | 0.65 | 0.24 | 0.22 |
| **Elastic Net Regression** | 1.57 | 0.65 | 0.05 | 0.03 |
| **Generalized Additive Model with Boosting** | 1.56 | 0.39 | -0.06 | -0.07 |
| **Gaussian Processes Regression Linear** | 1.71 | 1.07 | 0.14 | 0.18 |
| **Gaussian Processes Regression Polynomial** | 1.55 | 0.62 | 0.2 | 0.19 |
| **Gaussian Processes Regression Radial** | 1.65 | 0.48 | 0.22 | 0.19 |
| **Generalized Linear Model** | 1.72 | 1.1 | 0.14 | 0.19 |
| **Independent Component Regression** | 1.84 | 0.35 | 0.1 | 0.08 |
| **Kernel Partial Least Squares** | 1.53 | 0.63 | 0.19 | 0.18 |
| **Principal Component Regression** | 1.84 | 0.35 | 0.1 | 0.08 |
| **Random Forest** | 1.55 | -0.03 | -0.14 | -0.19 |
| **Quantile Regression with LASSO penalty** | 1.62 | 0.64 | 0.08 | 0.14 |
| **Relevance Vector Machine-Linear** | -11.09 | 0.38 | -12.95 | -0.06 |
| **Relevance Vector Machine-Polynomial** | 1.47 | 0.62 | 0.24 | 0.24 |
| **Relevance Vector Machine-Radial** | 1.62 | 0.84 | -0.03 | 0.05 |
| **Sparse Partial Least Squares** | 1.53 | 0.63 | 0.19 | 0.18 |
| **Support Vector Regression-Linear** | 1.61 | 1.05 | 0.25 | 0.34 |
| **Support Vector Regression -Polynomial** | 1.47 | 0.57 | 0.16 | 0.19 |
| **Support Vector Regression -Radial** | 1.45 | 0.56 | 0.24 | 0.26 |
| **Extreme Gradient Boosting** | 1.33 | 0.23 | 0.06 | 0.05 |
| **Females** | | | | |
| **Bayesian Generalized Linear Model** | 2.09 | 1.26 | -0.69 | -0.72 |
| **Bayesian Regularized Neural Network** | 1.19 | 0.44 | -0.04 | 0.05 |
| **Elastic Net Regression** | 0.96 | 0.22 | -0.37 | -0.3 |
| **Generalized Additive Model with Boosting** | 0.87 | -0.08 | -0.63 | -0.55 |
| **Gaussian Processes Regression Linear** | 1.91 | 1.08 | -0.41 | -0.42 |
| **Gaussian Processes Regression Polynomial** | 1.11 | 0.29 | -0.39 | -0.3 |
| **Gaussian Processes Regression Radial** | 1.12 | 0.07 | -0.29 | -0.19 |
| **Generalized Linear Model** | 2.1 | 10.15 | -0.7 | 2.52 |
| **Independent Component Regression** | 0.98 | -0.15 | -0.45 | -0.35 |
| **Kernel Partial Least Squares** | 1.23 | 0.48 | -0.2 | -0.11 |
| **Principal Component Regression** | 0.98 | -0.14 | -0.45 | -0.34 |
| **Random Forest** | 0.97 | -0.38 | -0.43 | -0.34 |
| **Quantile Regression with LASSO penalty** | 1.03 | 0.26 | -0.39 | -0.19 |
| **Relevance Vector Machine-Linear** | -11.67 | -0.2 | -13.17 | -0.33 |
| **Relevance Vector Machine-Polynomial** | 1.08 | 0.33 | -0.38 | -0.28 |
| **Relevance Vector Machine-Radial** | 1.07 | 0.41 | -0.66 | -0.45 |
| **Sparse Partial Least Squares** | 1.23 | 0.5 | -0.2 | -0.1 |
| **Support Vector Regression-Linear** | 1.74 | 1.15 | -0.29 | -0.11 |
| **Support Vector Regression -Polynomial** | 1.08 | 0.28 | -0.31 | -0.19 |
| **Support Vector Regression -Radial** | 1.05 | 0.21 | -0.27 | -0.16 |
| **Extreme Gradient Boosting** | 0.73 | -0.15 | -0.28 | -0.19 |
| ABCD: Adolescent Brain Cognitive Development; BrainAGE_corr_: Age-bias adjusted BrainAGE;; LASSO: least absolute shrinkage and selection operator; PING: Pediatric Imaging, Neurocognition, and Genetics Data Repository | | | | |

**Supplemental Figure 3.** **Correlations Between Predicted and Chronological Age (years) in the Cross-Validation Sets.**

The different algorithms are referenced by the function used for their implementation. Bayesian Generalized Linear Model (bayesglm); Bayesian Regularized Neural Network (brnn); Elastic Net Regression (enet);Generalized Additive Model with Boosting (gamboost);: Gaussian Processes Regression Linear (gaussprLinear); Gaussian Processes Regression Polynomial (gaussprPoly); Gaussian Processes Regression Radial (gaussprRadial); Generalized Linear Model (glm); Independent Component Regression (icr); Kernel Partial Least Squares (kernelpls); Principal Component Regression (pcr); Random Forest (rf); Quantile Regression with LASSO penalty (rqlasso); Relevance Vector Machine-Linear (rvmLinear3); Relevance Vector Machine-Polynomial (rvmPoly); rvmRadial: Relevance Vector Machine-Radial; Sparse Partial Least Squares (spls); Support Vector Regression-Linear (svmeLinear3); Support Vector Regression-Polynomial (svmPoly); Support Vector Regression-Radial (svmRadial); Extreme Gradient Boosting (xgbTree)

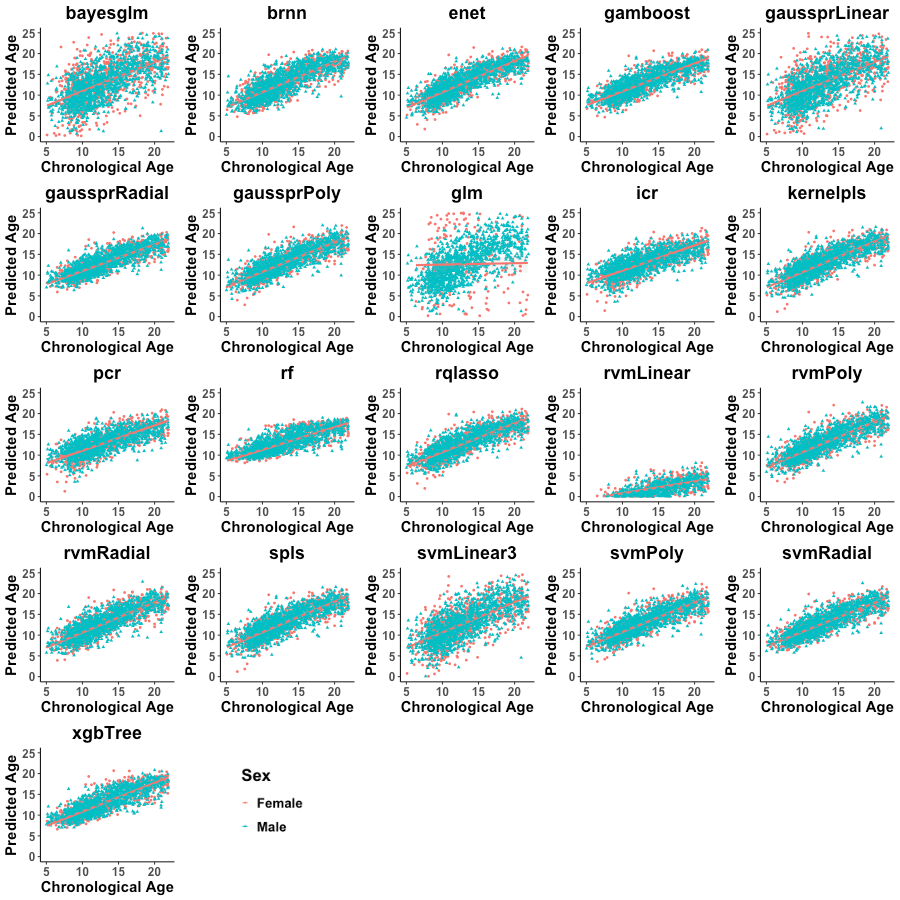

**Supplemental Figure 4.** **Correlations Between Brain Age Gap Estimation (BrainAGE) and Chronological Age (years) in the Cross-Validation Sets.**

The different algorithms are referenced by the function used for their implementation. Bayesian Generalized Linear Model (bayesglm); Bayesian Regularized Neural Network (brnn); Elastic Net Regression (enet);Generalized Additive Model with Boosting (gamboost);: Gaussian Processes Regression Linear (gaussprLinear); Gaussian Processes Regression Polynomial (gaussprPoly); Gaussian Processes Regression Radial (gaussprRadial); Generalized Linear Model (glm); Independent Component Regression (icr); Kernel Partial Least Squares (kernelpls); Principal Component Regression (pcr); Random Forest (rf); Quantile Regression with LASSO penalty (rqlasso); Relevance Vector Machine-Linear (rvmLinear3); Relevance Vector Machine-Polynomial (rvmPoly); rvmRadial: Relevance Vector Machine-Radial; Sparse Partial Least Squares (spls); Support Vector Regression-Linear (svmeLinear3); Support Vector Regression-Polynomial (svmPoly); Support Vector Regression-Radial (svmRadial); Extreme Gradient Boosting (xgbTree)

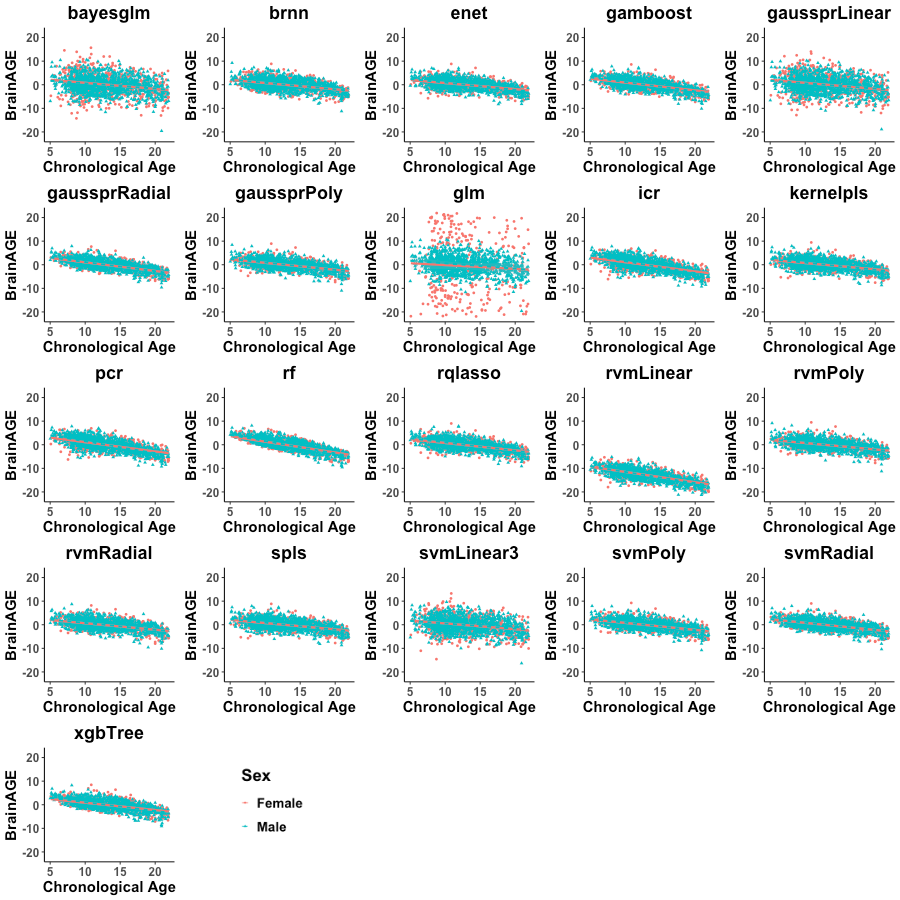

**Supplemental Figure 5.** **Correlations Between Age-Bias Corrected Brain Age Gap Estimation (BrainAge_corr_) and Chronological Age (years) in the Cross-Validation Sets.**

The different algorithms are referenced by the function used for their implementation. Bayesian Generalized Linear Model (bayesglm); Bayesian Regularized Neural Network (brnn); Elastic Net Regression (enet);Generalized Additive Model with Boosting (gamboost);: Gaussian Processes Regression Linear (gaussprLinear); Gaussian Processes Regression Polynomial (gaussprPoly); Gaussian Processes Regression Radial (gaussprRadial); Generalized Linear Model (glm); Independent Component Regression (icr); Kernel Partial Least Squares (kernelpls); Principal Component Regression (pcr); Random Forest (rf); Quantile Regression with LASSO penalty (rqlasso); Relevance Vector Machine-Linear (rvmLinear3); Relevance Vector Machine-Polynomial (rvmPoly); rvmRadial: Relevance Vector Machine-Radial; Sparse Partial Least Squares (spls); Support Vector Regression-Linear (svmeLinear3); Support Vector Regression-Polynomial (svmPoly); Support Vector Regression-Radial (svmRadial); Extreme Gradient Boosting (xgbTree)

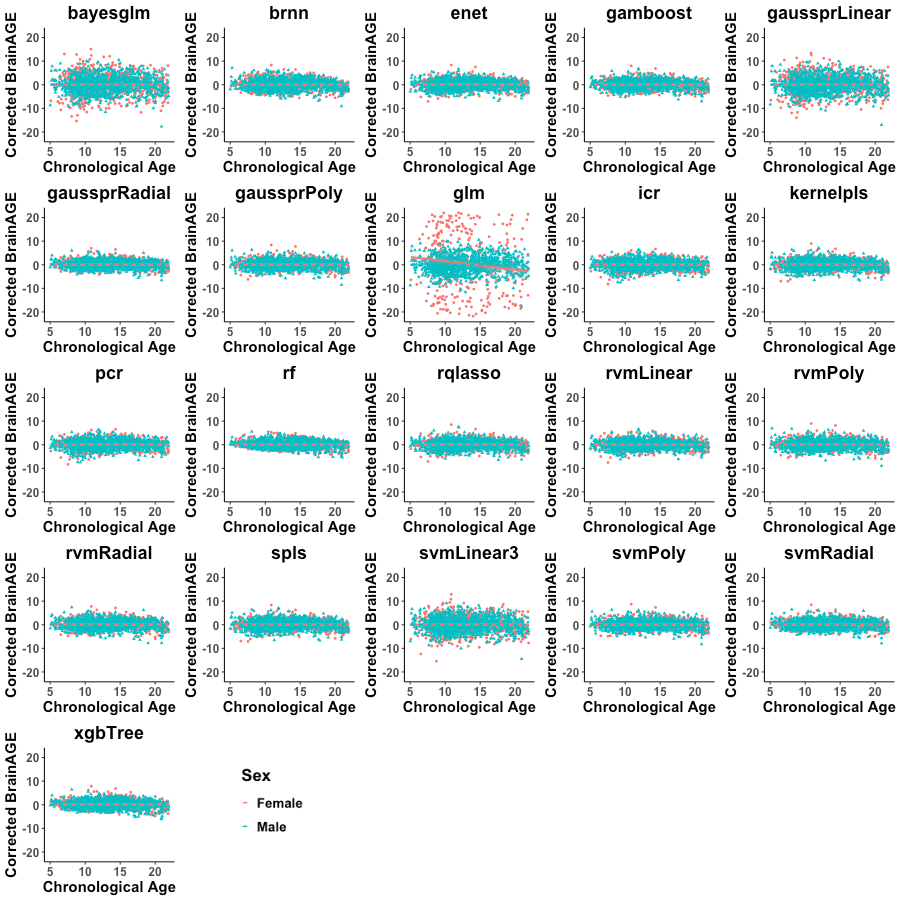

**Supplemental Figure 6.** **Correlations Between Brain Age Gap Estimation (BrainAGE) and Chronological Age (years) in the PING dataset.**

PING: Pediatric Imaging, Neurocognition, and Genetics Data Repository. The different algorithms are referenced by the function used for their implementation. Bayesian Generalized Linear Model (bayesglm); Bayesian Regularized Neural Network (brnn); Elastic Net Regression (enet);Generalized Additive Model with Boosting (gamboost);: Gaussian Processes Regression Linear (gaussprLinear); Gaussian Processes Regression Polynomial (gaussprPoly); Gaussian Processes Regression Radial (gaussprRadial); Generalized Linear Model (glm); Independent Component Regression (icr); Kernel Partial Least Squares (kernelpls); Principal Component Regression (pcr); Random Forest (rf); Quantile Regression with LASSO penalty (rqlasso); Relevance Vector Machine-Linear (rvmLinear3); Relevance Vector Machine-Polynomial (rvmPoly); rvmRadial: Relevance Vector Machine-Radial; Sparse Partial Least Squares (spls); Support Vector Regression-Linear (svmeLinear3); Support Vector Regression-Polynomial (svmPoly); Support Vector Regression-Radial (svmRadial); Extreme Gradient Boosting (xgbTree)

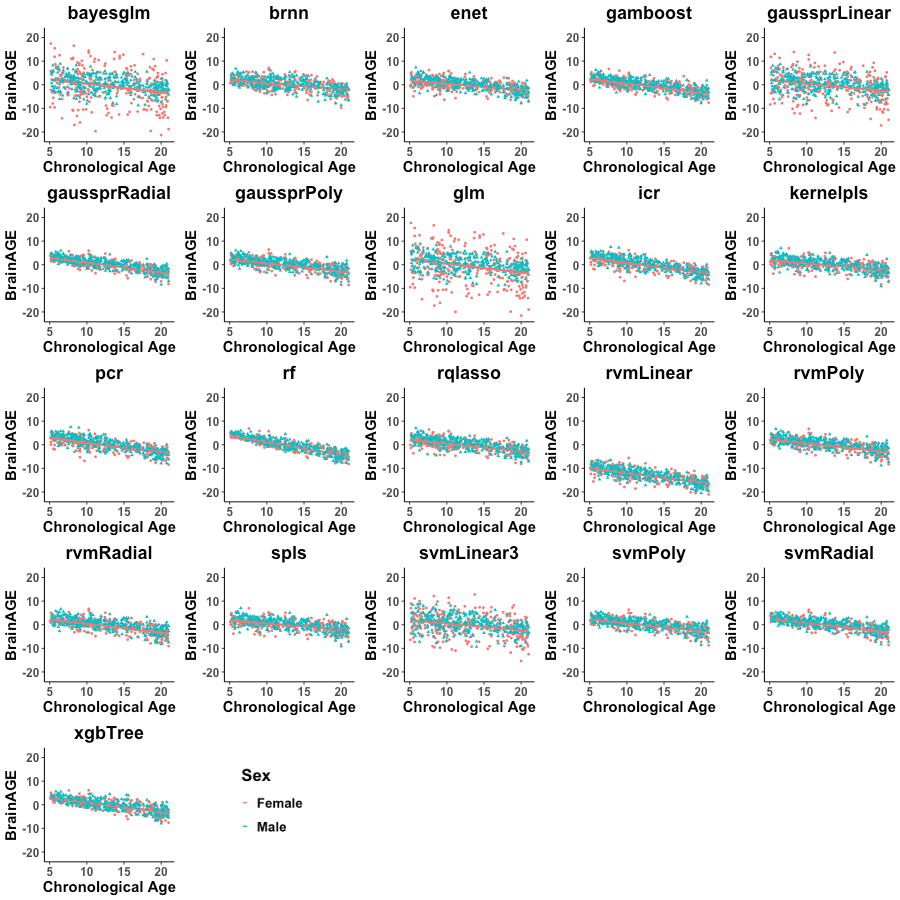

**Supplemental Figure 7.** **Correlations Between Age-Bias Corrected Brain Age Gap Estimation (BrainAGE_corr_) and Chronological Age (years) in the PING Dataset.**

PING: Pediatric Imaging, Neurocognition, and Genetics Data Repository. The different algorithms are referenced by the function used for their implementation. Bayesian Generalized Linear Model (bayesglm); Bayesian Regularized Neural Network (brnn); Elastic Net Regression (enet);Generalized Additive Model with Boosting (gamboost);: Gaussian Processes Regression Linear (gaussprLinear); Gaussian Processes Regression Polynomial (gaussprPoly); Gaussian Processes Regression Radial (gaussprRadial); Generalized Linear Model (glm); Independent Component Regression (icr); Kernel Partial Least Squares (kernelpls); Principal Component Regression (pcr); Random Forest (rf); Quantile Regression with LASSO penalty (rqlasso); Relevance Vector Machine-Linear (rvmLinear3); Relevance Vector Machine-Polynomial (rvmPoly); rvmRadial: Relevance Vector Machine-Radial; Sparse Partial Least Squares (spls); Support Vector Regression-Linear (svmeLinear3); Support Vector Regression-Polynomial (svmPoly); Support Vector Regression-Radial (svmRadial); Extreme Gradient Boosting (xgbTree)

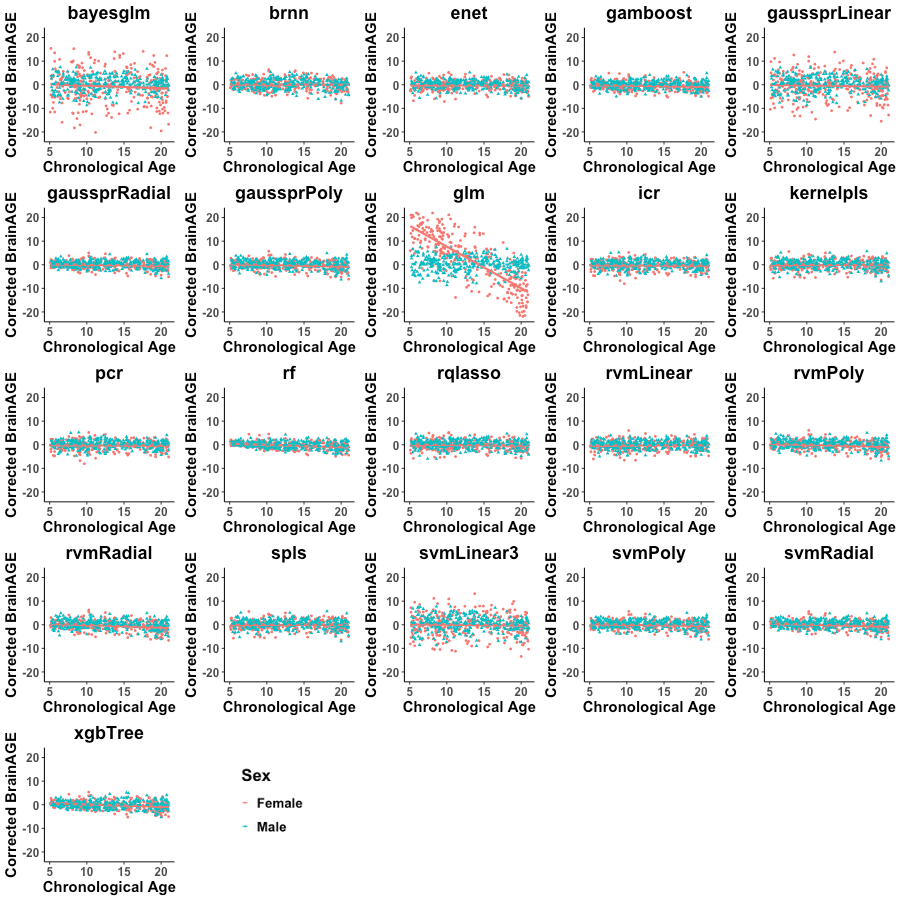

**Supplemental Figure 8.** **Age** **Bias-Corrected Mean Absolute Error (MAE_corr_) of the 21 Algorithms.** The figure presents the model performance in males and females in the hold-out test sets: the Adolescent Brain Cognitive Development (ABCD) study (Panel A) and the Pediatric Imaging, Neurocognition, and Genetics Data Repository (PING) (Panel B). The different algorithms are referenced by the function used for their implementation.

Bayesian Generalized Linear Model (bayesglm); Bayesian Regularized Neural Network (brnn); Elastic Net Regression (enet);Generalized Additive Model with Boosting (gamboost);: Gaussian Processes Regression Linear (gaussprLinear); Gaussian Processes Regression Polynomial (gaussprPoly); Gaussian Processes Regression Radial (gaussprRadial); Generalized Linear Model (glm); Independent Component Regression (icr); Kernel Partial Least Squares (kernelpls); Principal Component Regression (pcr); Random Forest (rf); Quantile Regression with LASSO penalty (rqlasso); Relevance Vector Machine-Linear (rvmLinear3); Relevance Vector Machine-Polynomial (rvmPoly); rvmRadial: Relevance Vector Machine-Radial; Sparse Partial Least Squares (spls); Support Vector Regression-Linear (svmeLinear3); Support Vector Regression-Polynomial (svmPoly); Support Vector Regression-Radial (svmRadial); Extreme Gradient Boosting (xgbTree)

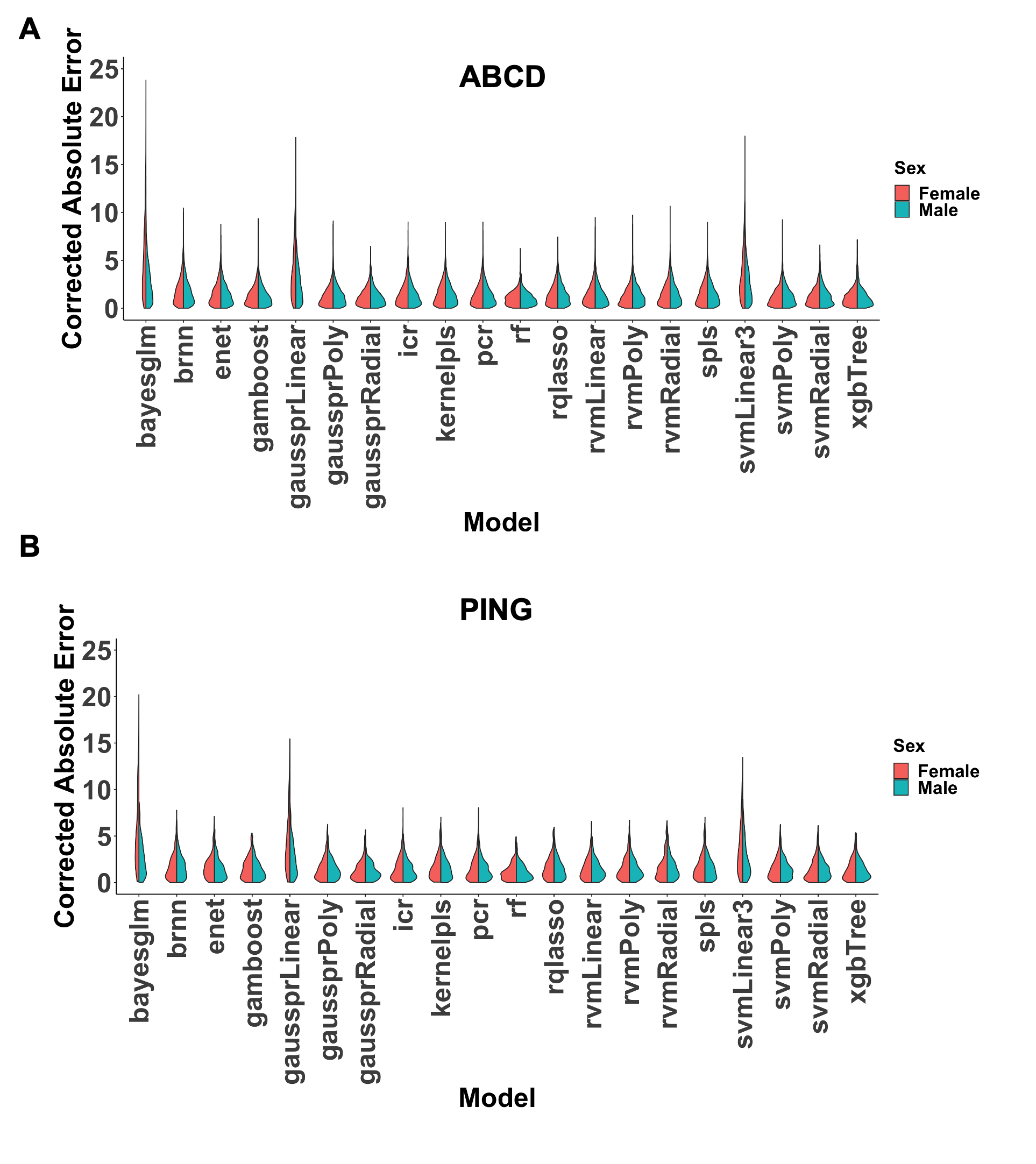

| **Supplemental Table 5.** **Top 20 Features Contributing to Age Prediction in the Three Best-Performing Algorithms by Sex** | | |
| --- | --- | --- |
| **Extreme Gradient Boosting** | **Random Forest** | **Support Vector Regression -Radial** |
| **Males** | | |
| Thick_rh_Cont_PFCl_3 | Thick_rh_Cont_PFCl_3 | Thick_lh_Limbic_OFC_3 |
| Thick_lh_Default_PCC_7 | Thick_lh_Default_PCC_7 | Area_rh_SomMot_33 |
| Thick_rh_Default_Temp_6 | Thick_lh_SalVentAttn_Med_3 | Thick_rh_SomMot_18 |
| Thick_lh_SalVentAttn_Med_3 | Thick_rh_Default_Temp_6 | Thick_lh_SomMot_26 |
| Thick_rh_DorsAttn_Post_5 | Thick_rh_DorsAttn_Post_5 | Area_rh_Limbic_OFC_5 |
| Thick_rh_SalVentAttn_Med_3 | Thick_lh_Vis_5 | Thick_rh_Default_PFCm_1 |
| Thick_lh_Limbic_OFC_5 | Thick_rh_SalVentAttn_Med_3 | Thick_lh_Vis_15 |
| Thick_rh_SalVentAttn_PFCl_1 | Thick_rh_SalVentAttn_PFCl_1 | Thick_lh_DorsAttn_Post_15 |
| Thick_lh_SomMot_16 | Thick_rh_Default_PFCm_1 | Area_rh_Cont_Par_2 |
| Thick_lh_SomMot_26 | Thick_lh_Limbic_OFC_5 | Thick_rh_Cont_PFCl_15 |
| Thick_rh_Cont_PFCl_5 | Thick_lh_Default_PFC_11 | Area_lh_SomMot_1 |
| Thick_lh_Default_PCC_6 | Thick_lh_Limbic_OFC_3 | Thick_rh_SomMot_38 |
| Thick_lh_Default_PCC_9 | Thick_rh_Vis_25 | Thick_lh_Default_Temp_6 |
| Thick_rh_SomMot_22 | Thick_lh_Vis_11 | Thick_rh_Cont_PFCl_6 |
| Thick_lh_Vis_22 | Thick_lh_Default_PCC_6 | Thick_rh_SomMot_22 |
| Thick_lh_Vis_11 | Thick_lh_Vis_22 | Area_lh_SomMot_26 |
| Thick_lh_Vis_5 | Thick_lh_Default_PCC_9 | Thick_lh_SomMot_16 |
| Thick_rh_SomMot_18 | Thick_lh_Default_PFC_4 | Area_rh_Limbic_TempPole_7 |
| Thick_lh_SalVentAttn_Med_5 | Thick_rh_Vis_24 | Area_rh_SomMot_27 |
| Thick_lh_Cont_PFCl_1 | Thick_lh_Vis_26 | Thick_lh_SalVentAttn_Med_5 |
| **Females** | | |
| Thick_rh_Cont_PFCl_3 | Thick_rh_Cont_PFCl_3 | Thick_lh_Default_PFC_3 |
| Thick_lh_SalVentAttn_Med_3 | Thick_lh_SalVentAttn_Med_3 | Thick_rh_Default_PFCm_1 |
| Thick_lh_Default_PFC_3 | Thick_lh_Default_PCC_7 | Thick_lh_SomMot_19 |
| Thick_lh_SalVentAttn_Med_5 | Thick_lh_Default_PFC_3 | Thick_rh_Default_PFCv_2 |
| Thick_rh_Default_Temp_6 | Thick_lh_SalVentAttn_Med_5 | Thick_lh_Cont_PFCl_5 |
| Thick_lh_Default_PCC_7 | Thick_lh_Default_PCC_6 | Thick_lh_SomMot_16 |
| Thick_lh_Default_PCC_6 | Thick_rh_Default_PFCm_3 | Thick_rh_Cont_PFCl_5 |
| Thick_lh_Default_PCC_9 | Thick_rh_Default_Temp_6 | Thick_lh_Vis_20 |
| Thick_lh_Vis_27 | Thick_lh_Vis_27 | Area_lh_SomMot_28 |
| Thick_rh_Default_PFCm_1 | Thick_lh_SalVentAttn_Med_2 | Thick_lh_Default_Temp_7 |
| Thick_rh_Default_PFCm_2 | Thick_rh_Default_PFCm_1 | Thick_lh_SalVentAttn_Med_2 |
| Thick_lh_DorsAttn_Post_5 | Thick_lh_DorsAttn_Post_5 | Thick_lh_SalVentAttn_Med_3 |
| Thick_lh_Vis_22 | Thick_rh_SalVentAttn_Med_3 | Thick_rh_SomMot_18 |
| Thick_lh_SalVentAttn_Med_2 | Thick_rh_SomMot_20 | Area_rh_Cont_PFCv_1 |
| Thick_lh_SomMot_19 | Thick_lh_Default_PCC_9 | Thick_rh_Default_Temp_6 |
| Thick_lh_SomMot_26 | Thick_lh_Default_PFC_9 | Area_lh_Limbic_OFC_1 |
| Thick_lh_Default_Temp_7 | Thick_rh_Cont_PFCl_5 | Area_rh_SomMot_33 |
| Thick_rh_SalVentAttn_Med_3 | Thick_rh_SalVentAttn_Med_1 | Thick_lh_Vis_22 |
| Thick_rh_Cont_PFCl_5 | Thick_lh_Limbic_OFC_5 | Thick_lh_Cont_PFCl_1 |
| Thick_lh_Cont_PFCl_5 | Thick_lh_Default_PFC_6 | Area_lh_SomMot_3 |
| Cont: Control Network; Default: Default Mode Network; DorsAttn: Dorsal Attention Network; lh: left hemisphere; Limbic: Limbic Network; rh: right hemisphere; SalVentAttn: Salience Ventral Attention Network; SomMot: Somatomotor; Vis: Visual Network. | | |

| **Supplemental Table 6.** **Mean absolute error (MAE) and Age-Bias Corrected Mean Absolute Error (MAE_corr_) of Models Trained in One Sex and Tested on the Other** | | | | |
| --- | --- | --- | --- | --- |
|  | **ABCD** | | | |
| **Algorithm (function name in caret package)** | **Models Trained on Females and Applied to Males** | | **Models Trained on Males and Applied to Females** | |
|  | **MAE** | **MAE_corr_** | **MAE** | **MAE_corr_** |
| **Extreme Gradient Boosting (xgbTree)** | 1.18 | 1.18 | 1.63 | 1.17 |
| **Random Forest Regression (rf)** | 1.25 | 1.02 | 1.63 | 1.15 |
| **Support Vector Regression - Radial Basis Function (svmRadial)** | 1.37 | 1.12 | 1.76 | 1.29 |
|  | **PING** | | | |
|  | **Models Trained in Females and Applied to Males** | | **Models Trained on Males and Applied to Females** | |
|  | **MAE** | **MAE_corr_** | **MAE** | **MAE_corr_** |
| **Extreme Gradient Boosting (xgbTree)** | 2.34 | 1.49 | 1.97 | 1.24 |
| **Random Forest Regression (rf)** | 2.78 | 1.33 | 2.55 | 1.13 |
| **Support Vector Regression - Radial Basis Function (svmRadial)** | 2.33 | 1.59 | 1.88 | 1.20 |
| ABCD: Adolescent Brain Cognitive Development; PING: Pediatric Imaging, Neurocognition, and Genetics Data Repository | | | | |

| **Supplemental Table 7. BrainAGE and BrainAGE_corr_ Derived from Models Trained on One Sex and Tested in the Other** | | | | |
| --- | --- | --- | --- | --- |
|  | **ABCD** | | | |
| **Algorithm (function name in caret package)** | **Models Trained on Females and Applied to Males** | | **Models Trained on Males and Applied to Females** | |
|  | **BrainAGE** | **BrainAGE_corr_** | **BrainAGE** | **BrainAGE_corr_** |
| **Extreme Gradient Boosting (xgbTree)** | 0.54 | -0.33 | 1.44 | 0.32 |
| **Random Forest Regression (rf)** | 1.02 | -0.31 | 1.51 | -0.09 |
| **Support Vector Regression – Radial Basis Function (svmRadial)** | 0.95 | 0.13 | 1.54 | 0.64 |
|  | **PING** | | | |
|  | **Models Trained on Females and Applied to Males** | | **Models Trained on Males and Applied to Females** | |
|  | **BrainAGE** | **BrainAGE_corr_** | **BrainAGE** | **BrainAGE_corr_** |
| **Extreme Gradient Boosting (xgbTree)** | -0.77 | -0.73 | 0.25 | 0.3 |
| **Random Forest Regression (rf)** | -0.73 | -0.7 | 0.03 | 0.05 |
| **Support Vector Regression - Radial Basis Function (svmRadial)** | -0.83 | -0.76 | 0.23 | 0.29 |
| ABCD: Adolescent Brain Cognitive Development; PING: Pediatric Imaging, Neurocognition, and Genetics Data Repository; BrainAGE_corr_: Age Bias-Corrected Brain-Age-Gap-Estimation | | | | |

**Supplemental Figure 9.** **ABCD Dataset: Mean Absolute Error per Parcellation Scheme in Each of the Three Best Performing Algorithms**

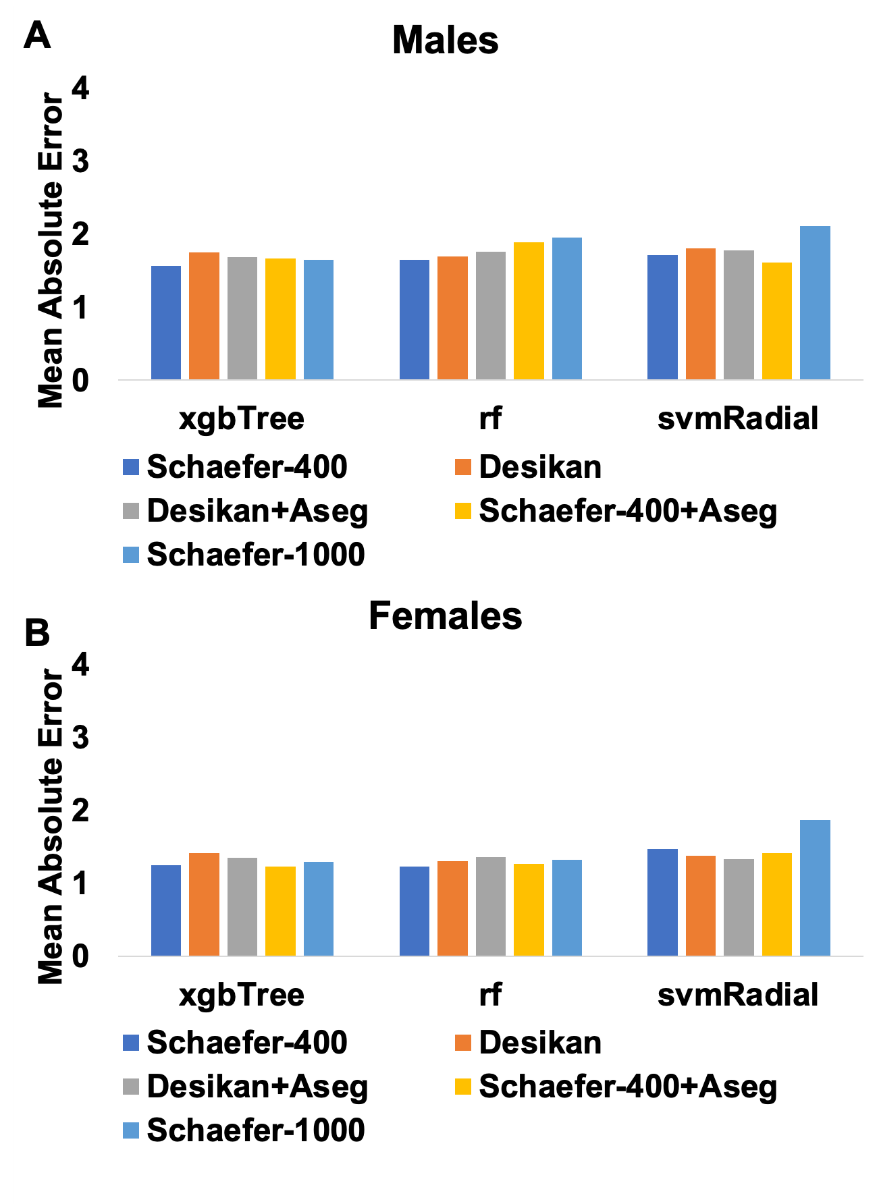

ABCD: Adolescent Brain Cognitive Development; Support Vector Regression - Radial Basis Function (svmRadial); Extreme Gradient Boosting (xgbTree); Random Forest Regression (rf)

**Supplemental Figure 10.** **PING Dataset:** **Mean Absolute Error per Parcellation Scheme in Each of the Three Best Performing Algorithms**

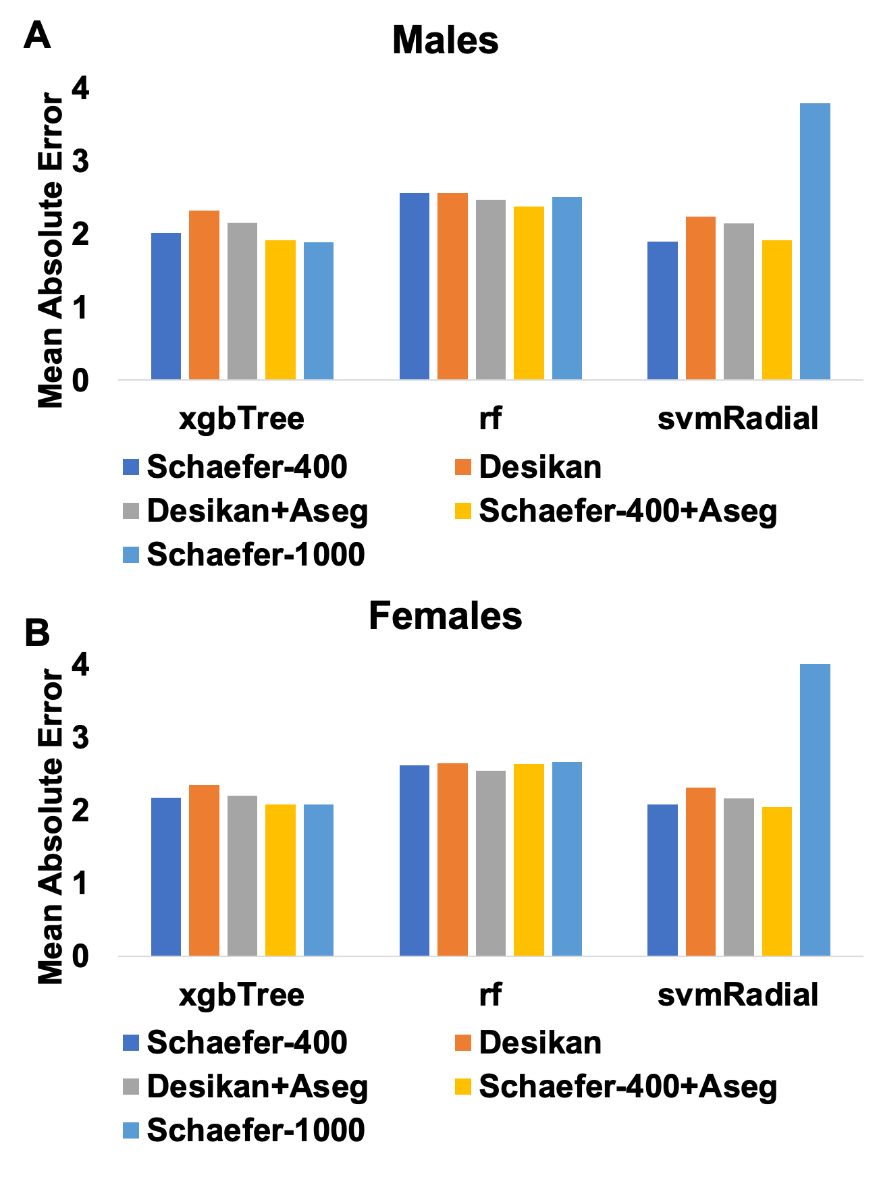

PING: Pediatric Imaging, Neurocognition, and Genetics Data Repository; Support Vector Regression - Radial Basis Function (svmRadial); Extreme Gradient Boosting (xgbTree); Random Forest Regression (rf)

**Supplemental Figure 11.** **Age-Bias Corrected Mean Absolute Error (MAE_corr_) of the Three Best Performing Algorithms as a Function of Sample Size**

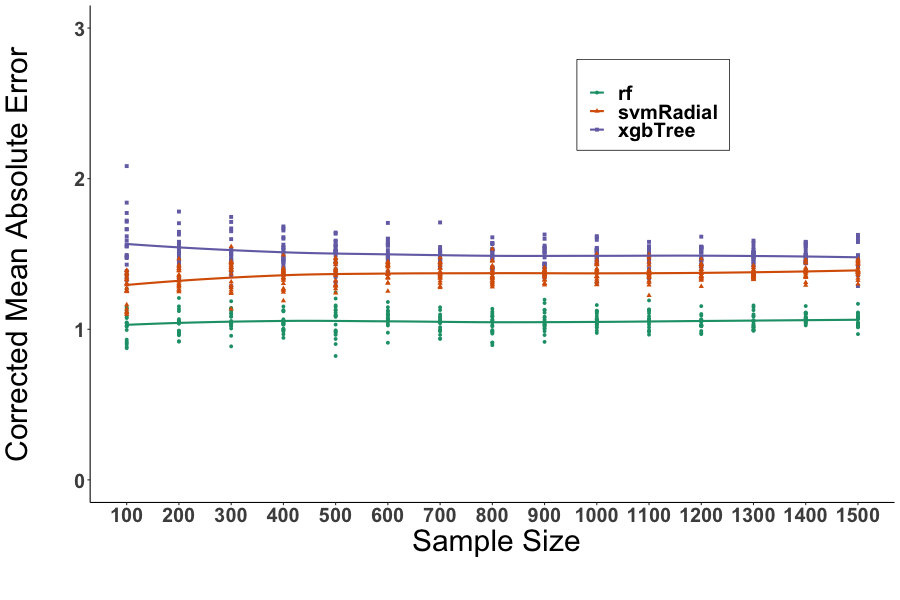

Data from the five datasets were randomly re-sampled without replacement to form sub-samples of 100-1500 participants at 100 increments; the process was repeated 20 times for each subsample. Support Vector Regression - Radial Basis Function (svmRadial); Extreme Gradient Boosting (xgbTree);

Random Forest Regression (rf)

| **Supplemental Table 8.** **Mean Absolute Error (MAE) and Age-Bias corrected Mean Absolute Error (MAE_corr_) Between 5 and 10 Repeats of the Tenfold Cross-Validation in the Three Best Performing Algorithms** | | | | |
| --- | --- | --- | --- | --- |
|  | **Five repeats of 5-fold cross-validation** | | **Ten repeats of 10-fold cross-validation** | |
|  | **ABCD** | | | |
| **Random Forest Regression (rf)** | **MAE** | **MAE_corr_** | **MAE** | **MAE_corr_** |
| Males | 1.65 | 1.09 | 1.66 | 1.1 |
| Females | 1.23 | 1.08 | 1.24 | 1.09 |
| **Support Vector Regression - Radial Basis Function (svmRadial)** | **MAE** | **MAE_corr_** | **MAE** | **MAE_corr_** |
| Males | 1.72 | 1.29 | 1.71 | 1.29 |
| Females | 1.47 | 1.21 | 1.46 | 1.24 |
| **Extreme Gradient Boosting (xgbTree)** | **MAE** | **MAE_corr_** | **MAE** | **MAE_corr_** |
| Males | 1.57 | 1.16 | 4.13 | 3.08 |
| Females | 1.25 | 1.19 | 5.71 | 4.87 |
|  | **PING** | | | |
| **Random Forest Regression (rf)** | **MAE** | **MAE_corr_** | **MAE** | **MAE_corr_** |
| Males | 2.57 | 1.14 | 2.57 | 1.13 |
| Females | 2.62 | 1.2 | 2.62 | 1.22 |
| **Support Vector Regression - Radial Basis Function (svmRadial)** | **MAE** | **MAE_corr_** | **MAE** | **MAE_corr_** |
| Males | 1.9 | 1.41 | 1.9 | 1.39 |
| Females | 2.08 | 1.38 | 2.09 | 1.39 |
| **Extreme Gradient Boosting (xgbTree)** | **MAE** | **MAE_corr_** | **MAE** | **MAE_corr_** |
| Males | 2.02 | 1.32 | 2.05 | 1.32 |
| Females | 2.17 | 1.41 | 2.23 | 1.37 |

| **Supplemental Table 9.** **Spearman's Correlation Coefficients Between the Number of Outliers, Mean Absolute Error (MAE) and Age-Bias Corrected Mean Absolute Error (MAE_corr_) for the Three Best Performing Algorithms** | | | | |
| --- | --- | --- | --- | --- |
| **Algorithm (function name in caret package)** | **Correlation Between Outliers and MAE** | | **Correlation Between Outliers MAE_corr_** | |
|  | **ABCD** | | | |
|  | **Males** | **Females** | **Males** | **Females** |
| **Extreme Gradient Boosting (xgbTree)** | 0.15 | 0.19 | 0.18 | 0.16 |
| **Random Forest Regression (rf)** | 0.24 | 0.25 | 0.13 | 0.13 |
| **Support Vector Regression - Radial Basis Function (svmRadial)** | 0.1 | 0.17 | 0.11 | 0.13 |
|  | **PING** | | | |
|  | **Males** | **Females** | **Males** | **Females** |
| **Extreme Gradient Boosting (xgbTree)** | -0.01 | 0.03 | -0.02 | 0.01 |
| **Random Forest Regression (rf)** | 0.08 | 0.12 | -0.01 | -0.02 |
| **Support Vector Regression - Radial Basis Function (svmRadial)** | -0.02 | 0.07 | 0.03 | -0.06 |
| ABCD: Adolescent Brain Cognitive Development; PING: Pediatric Imaging, Neurocognition, and Genetics Data Repository | | | | |

ADHD-200 Consortium, 2012. The ADHD-200 Consortium: A Model to Advance the Translational Potential of Neuroimaging in Clinical Neuroscience. Front Syst Neurosci 6, 62.

Di Martino, A., O'Connor, D., Chen, B., Alaerts, K., Anderson, J.S., Assaf, M., Balsters, J.H., Baxter, L., Beggiato, A., Bernaerts, S., Blanken, L.M., Bookheimer, S.Y., Braden, B.B., Byrge, L., Castellanos, F.X., Dapretto, M., Delorme, R., Fair, D.A., Fishman, I., Fitzgerald, J., Gallagher, L., Keehn, R.J., Kennedy, D.P., Lainhart, J.E., Luna, B., Mostofsky, S.H., Muller, R.A., Nebel, M.B., Nigg, J.T., O'Hearn, K., Solomon, M., Toro, R., Vaidya, C.J., Wenderoth, N., White, T., Craddock, R.C., Lord, C., Leventhal, B., Milham, M.P., 2017. Enhancing studies of the connectome in autism using the autism brain imaging data exchange II. Sci Data 4, 170010.

Di Martino, A., Yan, C.G., Li, Q., Denio, E., Castellanos, F.X., Alaerts, K., Anderson, J.S., Assaf, M., Bookheimer, S.Y., Dapretto, M., Deen, B., Delmonte, S., Dinstein, I., Ertl-Wagner, B., Fair, D.A., Gallagher, L., Kennedy, D.P., Keown, C.L., Keysers, C., Lainhart, J.E., Lord, C., Luna, B., Menon, V., Minshew, N.J., Monk, C.S., Mueller, S., Muller, R.A., Nebel, M.B., Nigg, J.T., O'Hearn, K., Pelphrey, K.A., Peltier, S.J., Rudie, J.D., Sunaert, S., Thioux, M., Tyszka, J.M., Uddin, L.Q., Verhoeven, J.S., Wenderoth, N., Wiggins, J.L., Mostofsky, S.H., Milham, M.P., 2014. The autism brain imaging data exchange: towards a large-scale evaluation of the intrinsic brain architecture in autism. Mol Psychiatry 19, 659-667.

Hagler, D.J., Jr., Hatton, S., Cornejo, M.D., Makowski, C., Fair, D.A., Dick, A.S., Sutherland, M.T., Casey, B.J., Barch, D.M., Harms, M.P., Watts, R., Bjork, J.M., Garavan, H.P., Hilmer, L., Pung, C.J., Sicat, C.S., Kuperman, J., Bartsch, H., Xue, F., Heitzeg, M.M., Laird, A.R., Trinh, T.T., Gonzalez, R., Tapert, S.F., Riedel, M.C., Squeglia, L.M., Hyde, L.W., Rosenberg, M.D., Earl, E.A., Howlett, K.D., Baker, F.C., Soules, M., Diaz, J., de Leon, O.R., Thompson, W.K., Neale, M.C., Herting, M., Sowell, E.R., Alvarez, R.P., Hawes, S.W., Sanchez, M., Bodurka, J., Breslin, F.J., Morris, A.S., Paulus, M.P., Simmons, W.K., Polimeni, J.R., van der Kouwe, A., Nencka, A.S., Gray, K.M., Pierpaoli, C., Matochik, J.A., Noronha, A., Aklin, W.M., Conway, K., Glantz, M., Hoffman, E., Little, R., Lopez, M., Pariyadath, V., Weiss, S.R., Wolff-Hughes, D.L., DelCarmen-Wiggins, R., Feldstein Ewing, S.W., Miranda-Dominguez, O., Nagel, B.J., Perrone, A.J., Sturgeon, D.T., Goldstone, A., Pfefferbaum, A., Pohl, K.M., Prouty, D., Uban, K., Bookheimer, S.Y., Dapretto, M., Galvan, A., Bagot, K., Giedd, J., Infante, M.A., Jacobus, J., Patrick, K., Shilling, P.D., Desikan, R., Li, Y., Sugrue, L., Banich, M.T., Friedman, N., Hewitt, J.K., Hopfer, C., Sakai, J., Tanabe, J., Cottler, L.B., Nixon, S.J., Chang, L., Cloak, C., Ernst, T., Reeves, G., Kennedy, D.N., Heeringa, S., Peltier, S., Schulenberg, J., Sripada, C., Zucker, R.A., Iacono, W.G., Luciana, M., Calabro, F.J., Clark, D.B., Lewis, D.A., Luna, B., Schirda, C., Brima, T., Foxe, J.J., Freedman, E.G., Mruzek, D.W., Mason, M.J., Huber, R., McGlade, E., Prescot, A., Renshaw, P.F., Yurgelun-Todd, D.A., Allgaier, N.A., Dumas, J.A., Ivanova, M., Potter, A., Florsheim, P., Larson, C., Lisdahl, K., Charness, M.E., Fuemmeler, B., Hettema, J.M., Maes, H.H., Steinberg, J., Anokhin, A.P., Glaser, P., Heath, A.C., Madden, P.A., Baskin-Sommers, A., Constable, R.T., Grant, S.J., Dowling, G.J., Brown, S.A., Jernigan, T.L., Dale, A.M., 2019. Image processing and analysis methods for the Adolescent Brain Cognitive Development Study. Neuroimage 202, 116091.

Harms, M.P., Somerville, L.H., Ances, B.M., Andersson, J., Barch, D.M., Bastiani, M., Bookheimer, S.Y., Brown, T.B., Buckner, R.L., Burgess, G.C., Coalson, T.S., Chappell, M.A., Dapretto, M., Douaud, G., Fischl, B., Glasser, M.F., Greve, D.N., Hodge, C., Jamison, K.W., Jbabdi, S., Kandala, S., Li, X., Mair, R.W., Mangia, S., Marcus, D., Mascali, D., Moeller, S., Nichols, T.E., Robinson, E.C., Salat, D.H., Smith, S.M., Sotiropoulos, S.N., Terpstra, M., Thomas, K.M., Tisdall, M.D., Ugurbil, K., van der Kouwe, A., Woods, R.P., Zollei, L., Van Essen, D.C., Yacoub, E., 2018. Extending the Human Connectome Project across ages: Imaging protocols for the Lifespan Development and Aging projects. Neuroimage 183, 972-984.

Jernigan, T.L., Brown, T.T., Hagler, D.J., Jr., Akshoomoff, N., Bartsch, H., Newman, E., Thompson, W.K., Bloss, C.S., Murray, S.S., Schork, N., Kennedy, D.N., Kuperman, J.M., McCabe, C., Chung, Y., Libiger, O., Maddox, M., Casey, B.J., Chang, L., Ernst, T.M., Frazier, J.A., Gruen, J.R., Sowell, E.R., Kenet, T., Kaufmann, W.E., Mostofsky, S., Amaral, D.G., Dale, A.M., Pediatric Imaging, N., Genetics, S., 2016. The Pediatric Imaging, Neurocognition, and Genetics (PING) Data Repository. Neuroimage 124, 1149-1154.
